# GhostiPy: an efficient signal processing and spectral analysis toolbox for large data

**DOI:** 10.1101/2021.04.30.442217

**Authors:** Joshua P. Chu, Caleb T. Kemere

## Abstract

Recent technological advances have enabled neural recordings consisting of hundreds to thousands of channels. As the pace of these developments continues to grow rapidly, it is imperative to have fast, flexible tools supporting the analysis of neural data gathered by such large scale modalities. Here we introduce `ghostipy` (**g**eneral **h**ub **o**f **s**pectral **t**echniques **i**n **Py**thon), a Python open source software toolbox implementing various signal processing and spectral analyses including optimal digital filters and time-frequency transforms. `ghostipy` prioritizes performance and efficiency by using parallelized, blocked algorithms. As a result, it is able to outperform commercial software in both time and space complexity for high channel count data and can handle out-of-core computation in a user-friendly manner. Overall, our software suite reduces frequently encountered bottlenecks in the experimental pipeline, and we believe this toolset will enhance both the portability and scalability of neural data analysis.

## Introduction

Advancements in neural recording technologies have enabled the collection of large data in both space (high density/channel count) and time (continuous recordings). During subsequent analysis, the scale of the data induces certain challenges which may manifest as the following scenarios: (1) analysis code takes a long time to complete (high time complexity), (2) code is unable to complete due to insufficient memory on the hardware (high spatial complexity). Moreover the scientist may have difficulty finding existing tools that address both (1) and (2) and that implement the desired analyses.

Although a potential remedy is to simply upgrade the hardware, it is not an acceptable solution for scientists desiring portability of analyses. In those situations hardware resources may be limited (e.g. using a laptop at the airport). We thus took an alternate approach by efficiently implementing analyses that would trivially scale for different hardware configurations. Our solution is `ghostipy` (**g**eneral **h**ub **o**f **s**pectral **t**echniques **i**n **Py**thon), a free and open source Python toolbox that attempts to optimize both time and space complexity in the context of spectral analyses. Methods include linear filtering, signal envelope extraction, and spectrogram estimation, according to best practices. `ghostipy` is designed for general purpose usage; while well suited for high density continuous neural data, it works with any arbitrary array-like data object.

In this paper we first describe `ghostipy`’s software design principles to increase efficiency. We then elaborate on featured methods along with code samples illustrating user friendliness of the software. Finally we benchmark our software against a comparable implementation, and we discuss strategies for working under an out-of-core (when data cannot fit into system memory) processing context.

## Materials and methods

An overview of implemented methods can be found in Table 1. Excluding out-of-core support, it is possible to use multiple different packages to achieve the same functionality. However the mix-and-match approach can reduce user friendliness since application programming interfaces (APIs) differ across packages and dependency management is more difficult. We believe our unified package provides an attractive solution to this challenge.

**Table 1.** Features implemented by `ghostipy` compared to existing software [27, 1, 28, 22, 13, 18, 12]

|  | Python | Overlap<br>save<br>convolu-<br>tion | Multi-<br>taper<br>method | Hilbert<br>Trans-<br>form | Morse<br>CWT | Synchro-<br>squeezed<br>trans-<br>form | Out-of-core |
| --- | --- | --- | --- | --- | --- | --- | --- |
| Ghostipy | + | + | + | + | + | + | + |
| SciPy | + | + | - | + | - | - | - |
| Chronux | - | - | + | - | - | - | - |
| Elephant | + | + | - | + | - | - | - |
| Brain-<br>storm | + | + | + | + | - | - | - |
| PyWt | + | - | - | - | - | + | - |
| Field<br>Trip | - | + | + | + | - | - | - |
| MNE | + | + | + | + | - | - | - |
| MATLAB | - | + | + | + | + | + | - |

### Software Design Considerations

As previously noted, successful completion of analyses may be hampered by long computation times or lack of system memory. Specifically, algorithmic time and space complexity are major determinants for the efficiency and performance of a software method. In general it is difficult to optimize both simultaneously. For example, time complexity may be reduced by increasing hardware parallelization, at the expense of higher space complexity (memory requirements). While we sought to lower both kinds of complexity compared to existing solutions, we gave space complexity a higher priority. Stated concretely, slow computation time is primarily a nuisance, but failure to complete an analysis due to insufficient memory is catastrophic.

Our design decision to prioritize space complexity was particularly critical because it directly influenced which backend library we chose for the Fast Fourier Transform (FFT), an operation used in the majority of `ghostipy`’s methods. While investigating the different options, we saw that `numpy` currently uses the `pocketfft` backend [26, 20]. When accelerated with Intel’s `MKL` library, it can be slightly faster than `FFTW` [6]. However, we have found `FFTW` [9, 10] superior for memory management and better suited for arbitrary length FFTs, including prime and odd numbers. An additional benefit of `FFTW` was its multithreaded capabilities. We therefore selected `FFTW` as our FFT backend.

To lower space complexity we used blocked algorithms, including overlap save convolution, which is not offered in any of the standard Python numerical computing libraries such as `numpy` or `scipy` [26, 27]. This approach enabled us to process very large data that could not fit in memory (also known as out-of-core processing).

Throughout our code, we also employed other strategies such as in-place operations. To lower the time complexity, we used efficient lengths of FFTs wherever possible, and we leveraged modern computing hardware by parallelizing our algorithms. For example a wavelet transform can be trivially parallelized since the transform for each scale is not dependent on other scales.

### Multitaper Method

Users often wish to perform a spectral decomposition on a signal of interest. This can be accomplished by using the multitaper method [25, 19]. The technique is well-suited to reduce the variance of a spectrum estimate, which is particularly useful when working with noisy neural data. The spectrum estimate is obtained as an average of multiple statistically independent spectrum estimators for a discrete signal *x*[*n*] with sampling frequency *f*_*s*_:

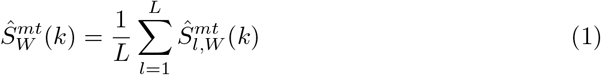

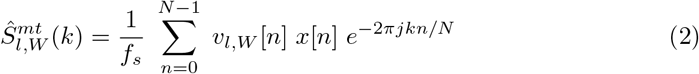

Given the length of data *N* and a smoothing half-bandwidth *W*, the tapers *v*_*l,W*_ [*n*] are computed by solving for vectors that satisfy the energy and orthogonality properties

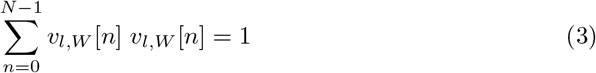

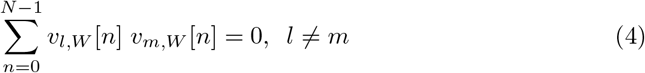

An example using `ghostipy` is shown in Figure 1.

**Fig 1.**
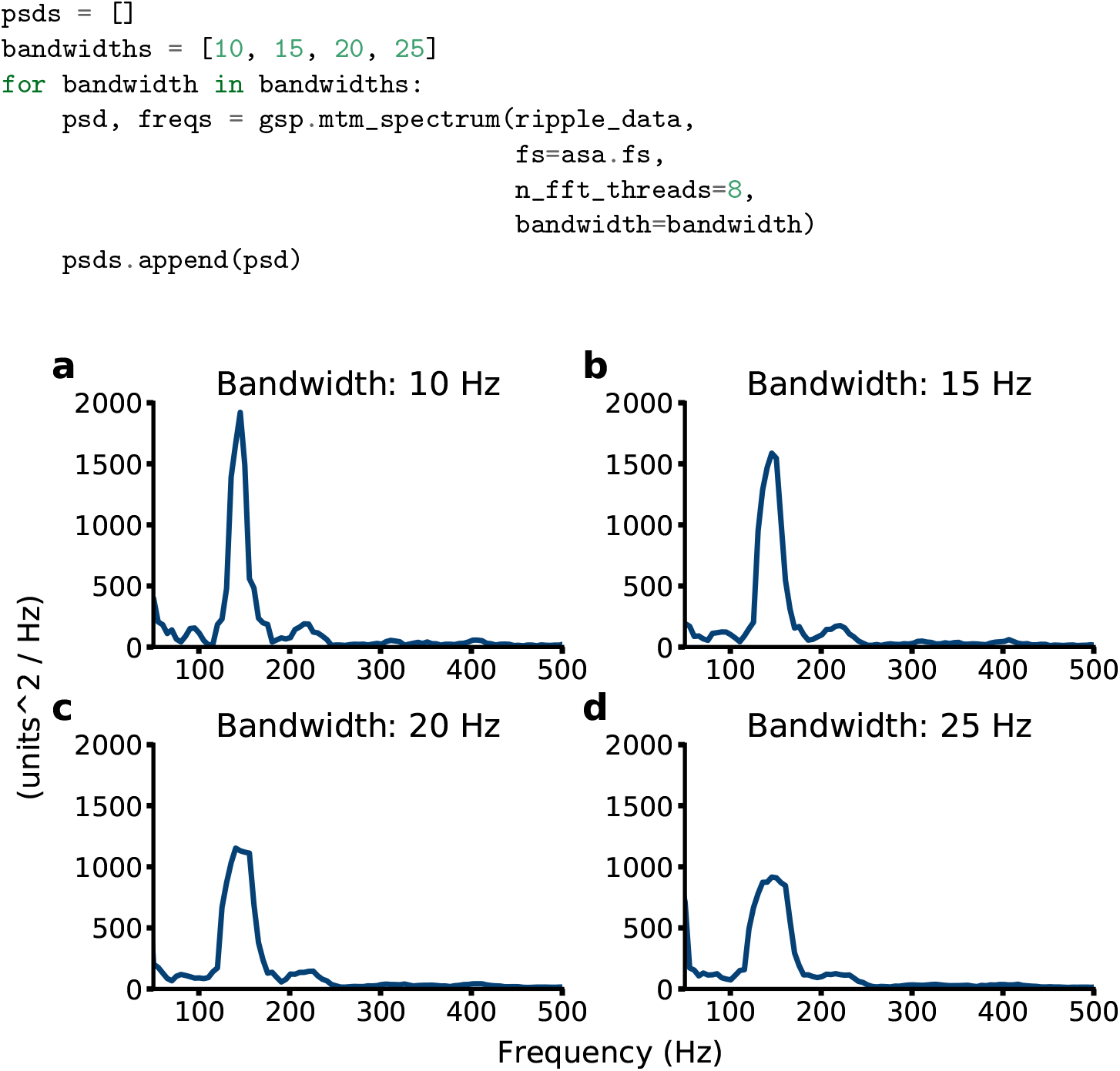
Mutitaper spectrum estimates can be readily generated. The example uses a sharp wave ripple event, where energy occurs mainly between 100 and 250 Hz.

### Continuous Wavelet Transform

Neuroscientists often use a continuous wavelet transform (CWT) to study transient oscillatory activity. The CWT itself is defined in the time domain by

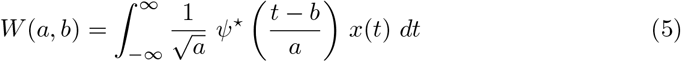

where *ψ*(…) is the mother wavelet function. The transform represents a two-dimensional decomposition in the scale (*a*) and time (*b*) planes. In the frequency domain, the CWT is given by the inverse Fourier transform of

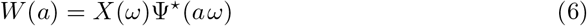

for a given scale (*a*), where *X* and Ψ are the Fourier transforms of *x* and *ψ*, respectively.

Many mother wavelet functions have been investigated in the literature, but we have focused on the analytic wavelets, as they are found to be superior, particularly for estimating phase [17, 14, 16, 15]. We have implemented the analytic Morse, Morlet, and Bump wavelets, whose respective frequency domain definitions are

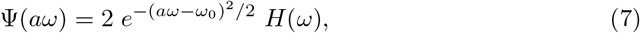

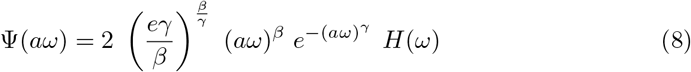

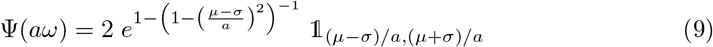

where 𝟙_(*µ*−*σ*)*/a*,(*µ*+*σ*)*/a*_ is the indicator function for the interval (*µ* − *σ*)*/a* ≤ *ω* ≤ (*µ* + *σ*)*/a* and *H*(*ω*) is the Heaviside step function. In our implementation, we use Equation 6 to compute the CWT.

Note that in practice the timeseries *x*(*t*) is sampled, and the CWT is likewise sampled. Then equation 6 becomes a pointwise complex multiplication of discrete Fourier transforms, where the discretized angular frequencies *ω*_*k*_ are determined by

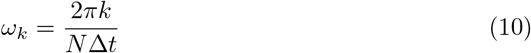

where *N* is the number of data samples and Δ*t* is the sampling interval.

For electrophysiological data, a typical wavelet analysis will require computing Equation 6 for 50-500 scales. This is an obvious candidate for parallelization since the wavelet transform for each scale can be computed independently of the others. We use a backend powered by `dask` to carry out the parallelization [21]. Users can set the number of parallel computations to execute and thereby leverage the multicore capabilities offered by modern computing hardware.

### Synchrosqueezing Transform

One disadvantage of the wavelet transform is that its frequency resolution decreases as the temporal resolution increases. Strictly speaking the CWT results in information contained in the (time, scale) plane, but a single frequency is typically assigned to each scale. Regardless, spectral smearing can be observed for at higher frequencies/lower scales. However, [4, 24] showed the synchrosqueezing transform (SST) could mitigate this issue by transferring a CWT’s (time, scale) plane information to the (time, frequency) plane.

The synchrosqueezing transform proceeds as the following. For every scale *a*:

1. Compute the CWT *W* (*a*) using equation 6
2. Compute the partial derivative

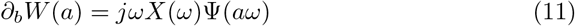
3. Compute the phase transform

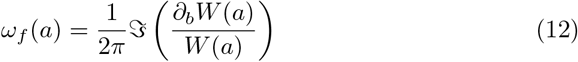

The phase transform contains the real frequencies each point in the CWT matrix should be assigned to. In practice the real frequency space is discretized, so the CWT points are assigned to frequency bins. Note that multiple CWT points at a given time coordinate *b* may map to the same frequency bin. In this situation, a given frequency bin is a simple additive accumulation of CWT points.

Note the similarity of the SST to the spectral reassignment algorithms in [11, 8]. However an important distinction is that the SST only operates along the scale dimension. In addition to preserving the temporal resolution of the CWT, this makes SST data easy to work with since uniform sampling can be maintained.

Overall, the spectrogram methods implemented by `ghostipy` give an experimenter a more complete picture of the time-varying spectral content of neural data. Figure 2 illustrates using the `scipy` standard spectrogram method along with `ghostipy`’s methods.

**Fig 2.**
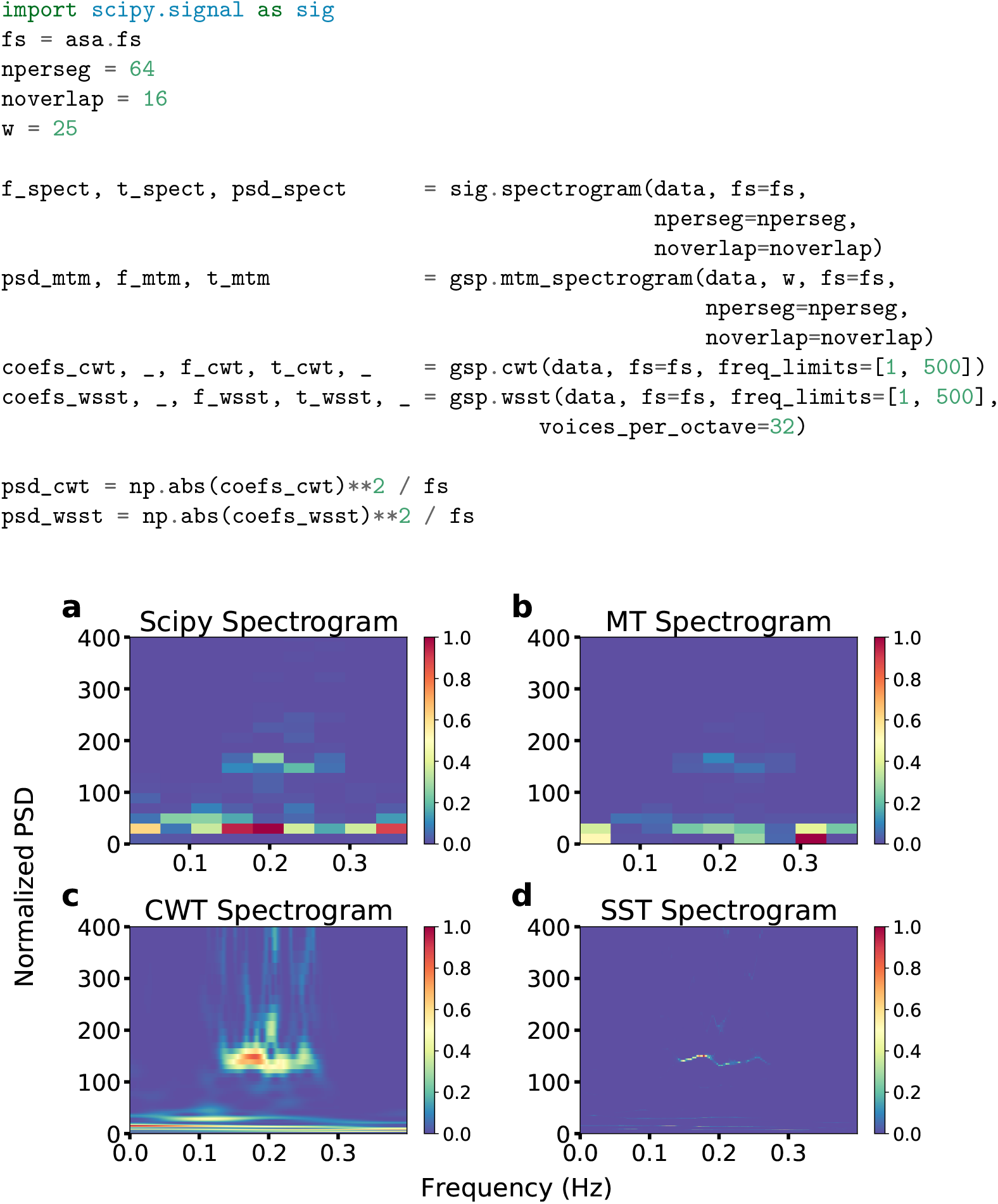
Users can leverage scipy’s spectrogram along with ghostipy’s methods to determine different time-frequency representations. The synchrosqueezed transform in (d) gives the overall sharpest time and frequency resolution.

### FIR Filter Design

In addition to the time-frequency transforms described above, `ghostipy` provides classical signal processing capabilities such as filtering data, using the efficient overlap save convolution. Filtering data is an ubiquitous operation, but before this stage, the filter must itself be designed. While this step may appear somewhat trivial, it can make a significant difference. Consider theta-gamma phase amplitude coupling (PAC), a phenomenon in which the amplitude of the gamma band (approximately 30-120 Hz) oscillation is modulated by the phase of the theta band (approximately 6-10 Hz) oscillation. [7] states that “It is not possible to measure significant amounts of PAC with amplitudes filtered with narrower bandwidths than the frequency of the modulatory rhythm”, but one of the early studies documenting theta-gamma coupling used 4 Hz bandwidth filters [3]. How could theta-gamma PAC have been discovered if the filters were nominally 4 Hz in bandwidth? The answer was that the software library used by the seminal paper scaled the transition bandwidth according to the passband center frequency, so the effective bandwidth was larger than the initially requested 4 Hz [7]. Theta-gamma coupling was thus a serendipitous discovery. Clearly it is important to control the transition bands of the filter, not just the passband. Existing packages such as `scipy` and `MNE` offer a variety of FIR filter design methods [27, 12]. However they suffer from certain issues, as documented below:

1. Least squares method: A solution may result in a filter with magnitude response effectively zero throughout. This situation is more common when designing filters with passband relatively low compared to the sampling rate.
2. Remez exchange method: The algorithm may simply fail to converge.
3. Window method: The transition bands cannot be controlled exactly, and optimality can not be defined, as is the case for the least squares (L2 optimal) and Remez exchange (L1 optimal)

Therefore `ghostipy`’s filter design uses the method defined in [2] for the following reasons:

1. It is simple to design. The computational complexity is similar to that of a window method and can be implemented on embedded hardware if desired.
2. Optimality can be defined, as it is optimal in the L2 sense.
3. Transition bands can be defined exactly, and the steepness of the passband rolloff can be controlled by the spline power parameter.
4. The filter impulse response can be defined analytically. Consequently its computation does not suffer from the failure modes of the least squares or Remez exchange methods, as those must solve systems of linear equations. In other words, the design process is reliable and stable.

This method designs a low pass filter according to

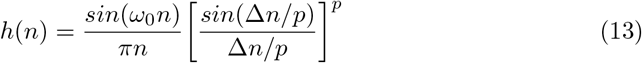

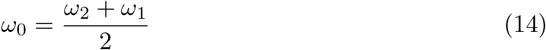

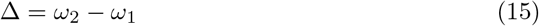

where *ω*_1_ and *ω*_2_ are radian frequencies defining the transition band boundaries.

`ghostipy` uses the low pass filter defined in 13 as a prototype to design more complicated filters. As a result, users can request filters with arbitrary magnitude response. Two examples are shown in Figure 3.

**Fig 3.**
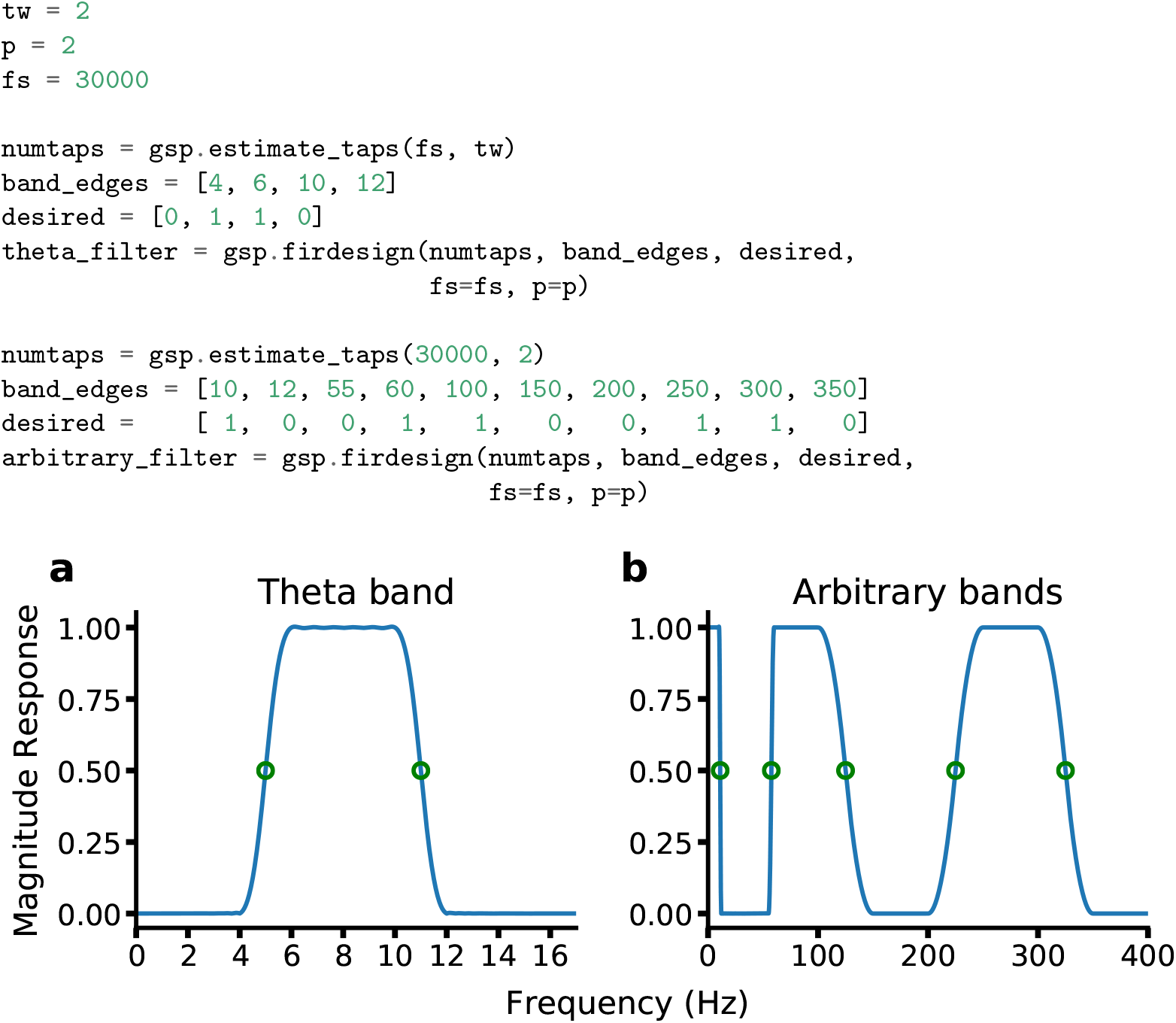
(a) A theta band filter designed for full bandwidth data. The specification of the transition bands allows for easy determination of critical frequencies. The −6 dB points are exactly the midpoints of the transition bands. (b) Filters with arbitrary pass and stop bands may also be designed.

## Results

Spectral analysis is ubiquitous in systems neuroscience experiments involving electrophysiology. One of the primary goals of our development was to facilitate clear and simple workflows with high performance. Of special note is that our tool enables wavelet-based spectral analysis using the Morse wavelet [15], a feature that has previously only been conveniently available in commercial tools such as Matlab. In particular, the Morse wavelet is exactly analytic and nicely parameterized to tradeoff frequency and temporal resolution. For very long data, most investigators will not be interested in the sub-Hz frequency components, and performance can be improved by restricting the frequencies/scales that are actually used for the transform.

A naive implementation of the Morse wavelet transform calculates untruncated wavelets the same length as the input data. This is often inefficient because it is equivalent to convolving the data with a time-domain wavelet mainly consisting of leading and trailing zeros. In our approach we exploit the fact that wavelets are finite in time and frequency, and we use an overlap-save algorithm to compute the CWT purely in the frequency domain. Note that the latter point is particularly critical: Due to the Gibbs phenomenon, using any time-domain representation of the wavelet may violate numerical analyticity for wavelet center frequencies near the Nyquist frequency. It is therefore necessary to use only the frequency domain representation of the wavelet. While we offer both traditional/naive and blockwise convolution implementations, the latter will give superior performance for longer-duration data. We believe that this is a valuable option for researchers and that this is the first tool which uses blockwise convolution to implement the CWT.

An example spectrogram of local field potentials recorded in area CA1 of the rat hippocampus is shown in figure 4. Clearly apparent are the theta oscillation, theta-nested gamma oscillations, and a sharp-wave-ripple, which occurs after the animal has stopped moving.

**Fig 4.**
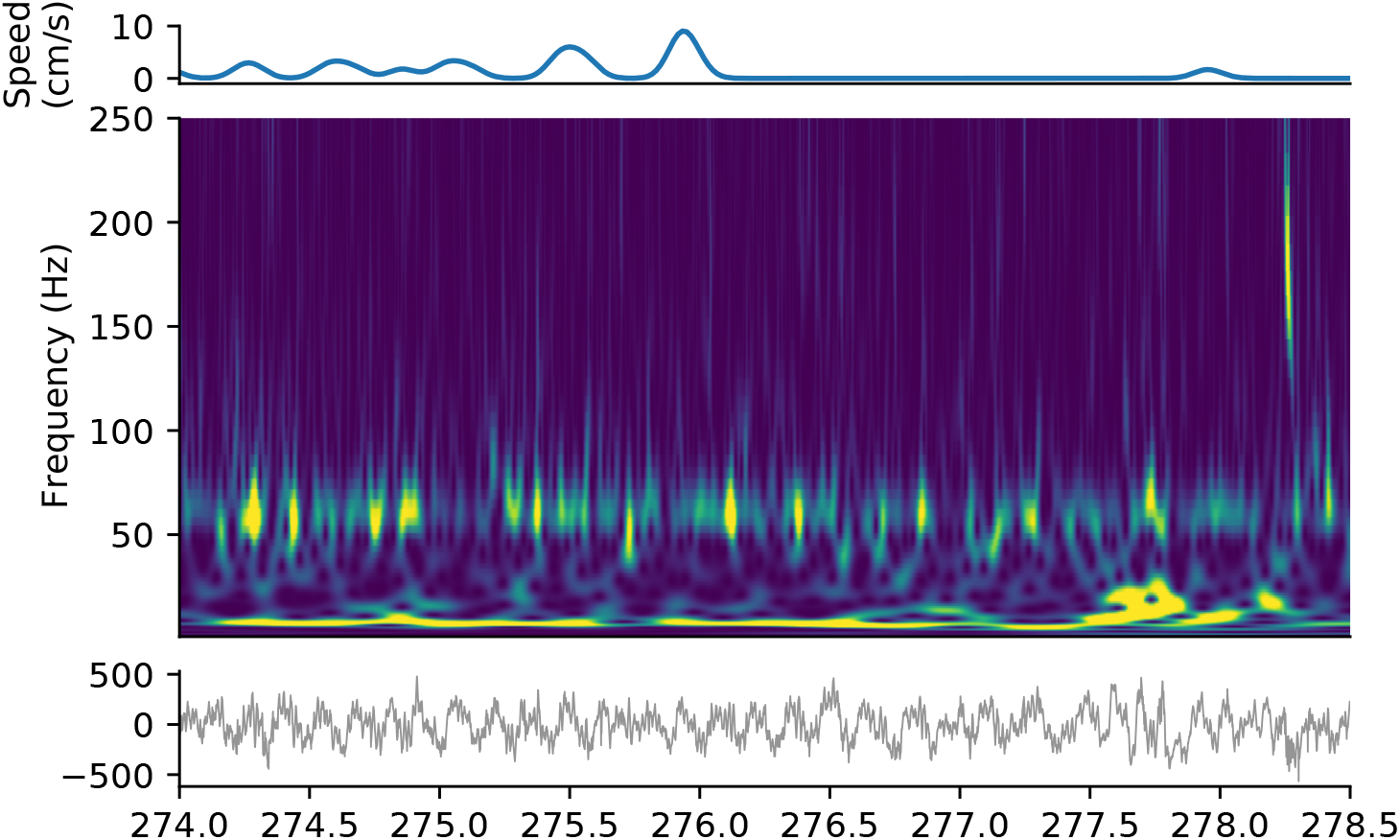
Spectrogram of local field potential recordings from area CA1 of the hippocampus of a rat during exploration (middle), with movement speed (top) and the raw electrophysiological signal (bottom). A number of features of the hippocampal rhythms can be noted in this example, including the pervasive theta oscillation (~8 Hz), theta-nested gamma oscillations (~60 Hz) during movement, and, towards the end, a sharp wave ripple (~200 Hz). An example notebook replicating this figure can be found [online](https://colab.research.google.com/drive/1TnNtEGQXEjfaxZTqtgexf4-S5BOIb6eu?usp=sharing).

### Performance and Complexity

The calculation of the CWT is computationally intensive and consequently a good method to benchmark performance. Of the software packages listed in Table 1, only `MATLAB` offered an equivalent solution. It was thus chosen as the reference to compare our implementation against. Figure 5 shows that our implementation results in faster computation times and better memory usage.

**Fig 5.**
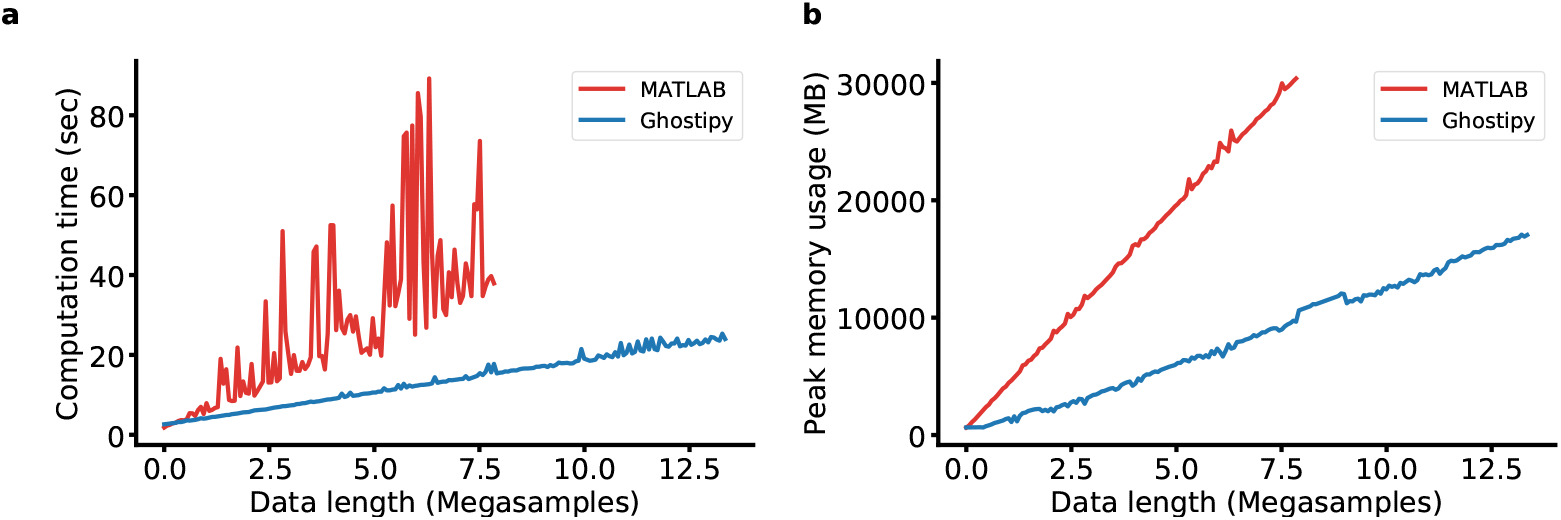
Our implementation of the Morse continuous wavelet transform outperforms `MATLAB`’s in both (a) time and (b) space complexity. Note that `MATLAB` was unable to complete execution for the full range of the test parameter (data length) due to out-of-memory exceptions. The test machine was an Intel Core i7-4790 (8 hyperthreads) equipped with 32 GB RAM.

It is not entirely clear what accounts for the higher jaggedness in the `MATLAB` curves from Figure 5. A possible explanation is that the FFT computation is less efficient for an odd-length transform, but the magnitude of the spikes in the curve is surprising given that `MATLAB`’s FFT backend also uses `FFTW`. Regardless, we have demonstrated that our implementation is able to achieve lower time and space complexity. When using the functionality offered by `ghostipy`, three primary scenarios arise with regards to the sizes of data involved in the processing:

1. both the input and output data fit into core memory
2. the input fits into core memory but the output does not
3. neither the input nor the output fit

In all of the previous examples, we have restricted ourselves to case 1. However, with the ever-increasing sizes of data, the other cases will inevitably be encountered. Case 2 may arise when attempting to generate spectrograms. As the input is a single channel, memory constraints are rarely an issue. For example, even a 10 hr LFP recording sampled at 1 kHz and saved as 64-bit floating point values will require less than 300 MiB of memory. However, the size of a wavelet spectrogram computed from this data will be directly proportional to the number of scales/frequencies. For a typical range of 1 to 350 Hz at 10 voices per octave, this amount to a space requirement of 85 times that of the input data. Given that this can well exceed the core memory size of a machine, `ghostipy`’s CWT routine can also accept a pre-allocated output array that is stored on disk (Figure 6).

**Fig 6.**
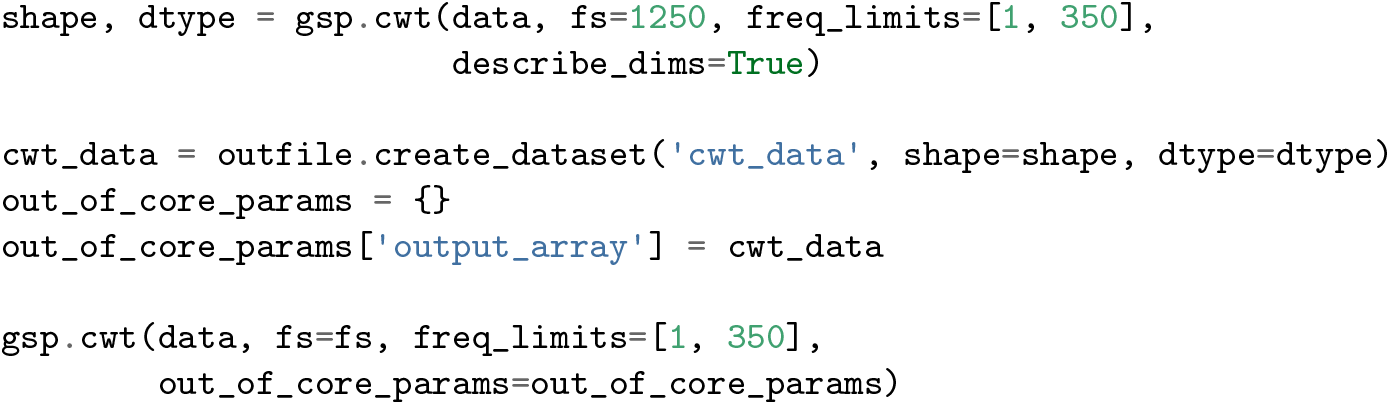
Example code when the output array is too large for main memory. The CWT method is first executed as a dry run to compute the necessary array sizes.

Case 3 may arise when a user wishes to filter many channels of full bandwidth data. One use case is a 1 hour recording for a 256 channel probe sampled at 30 kHz and stored as a 2-byte signed integer type; already this requires 51 GiB. Our strategy is similar to case 2, where an output array is allocated and stored on disk. As for the input, it is read in chunks, and the size of these can be chosen to lower memory usage, although potentially at a cost to computation time. The code in Figure 7 illustrates an example:

**Fig 7.**
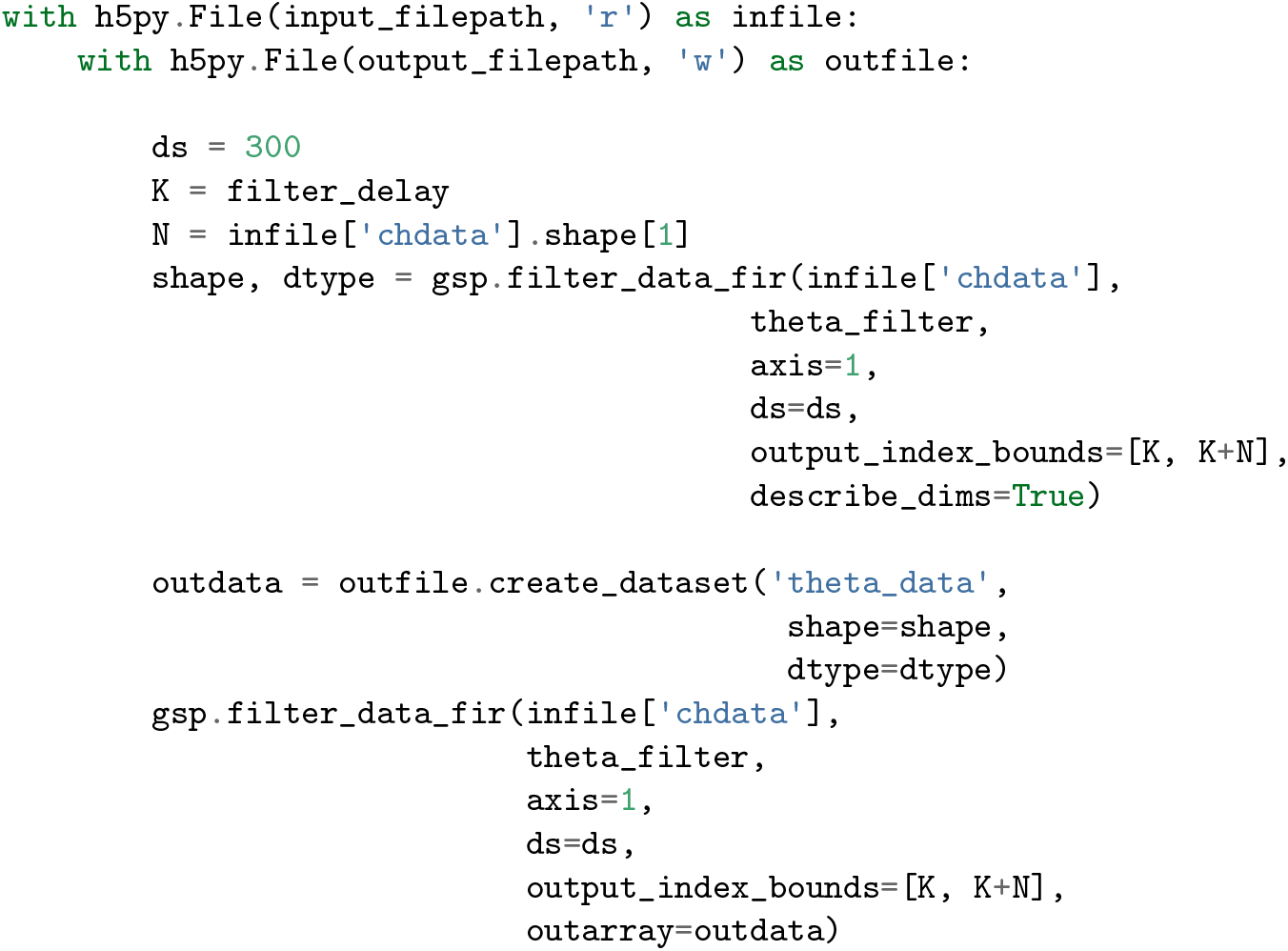
Filtering data from a large array stored on disk and likewise storing the output on disk. Similar to the CWT out-of-core features, the method is called once as a dry run to compute array sizes, which the user can then pass in to store the result. The filtering method also allows to correct for the delay of the filter and to downsample without storing any intermediate results. Although example uses the `h5py` library, any object that behaves like an array can be used.

Several points can be made about the scheme in Figure 7. Our method allows for downsampling during the convolution, which can reduce the number of stages in a computational scheme. Given full bandwidth data, a traditional strategy to filter to the theta band would look like the following:

1. Apply an anti-aliasing filter.
2. Downsample to obtain LFP.
3. Store the LFP to disk.
4. Apply a theta-band filter.
5. Downsample this output.
6. Save the result.

Using `ghostipy`’s method, it is not necessary to generate the intermediate LFP. To our knowledge, we do not know of other software that allows out-of-core filtering and downsampling in a single function call. The result is a simultaneous reduction in time and space complexity, by storing only the downsampled result and by filtering only once. Filtering to the theta band is now simplified to the following steps:

1. Apply a theta filter to the full bandwidth data.
2. Downsample the result.
3. Save the result to disk.

## Conclusion

We have described the key features of `ghostipy` and given examples of its ease of use to perform computations efficiently. Users can thus conduct exploratory spectral analyses quickly across a range of parameters while reducing their concerns for running out of memory, especially since out-of-core computation is supported for many of the methods. Thus we believe `ghostipy` is well-suited to handle the ever-increasing size of experimental data.

In the future we plan to improve `ghostipy` with various enhancements. For example, currently the methods are designed to offer the user a lot of low level control over areas such as multithreading, and to work with raw array types. However, users may desire a higher-level API. For this reason we believe it would be a worthwhile endeavor to incorporate our work into frameworks such as `NWB` [23]; this would also facilitate more widespread adoption. Lastly, there are other analyses we could implement, including the adaptive multitaper method [19] and other time-frequency reassignment techniques similar to the synchrosqueezing transform [5].

Our primary contribution is improving the ease and speed at which data analysis can be conducted, by developing user-friendly software implementing efficient algorithms well-suited for large data sizes. This point is specifically demonstrated by our ability to outperform existing solutions in space and time complexity, and to run computations even in out-of-core memory conditions, which enables machines with 1-10s of GBs of memory to process data on the scale of 10-100s GBs and higher. In these ways, we have increased the accessibility of neural data analysis by enabling it to be run on hardware such as laptops, a scenario that often was not previously possible.

Lastly, the software we developed has a much larger potential impact than the scope described in this paper. Although many of the examples given in this paper were specific to extracellular rodent hippocampal data, the functionality we implemented is intentionally generic and applicable to many fields. As an example, our code can easily be adapted for use in real-time processing, whether running on embedded hardware or on a laptop in a clinical EEG setting. Given the functionality already developed and the full scope of our work, we are optimistic that `ghostipy` can help accelerate modern scientific progress.

## Supporting information

Notebooks for figures

## Acknowledgements

We thank Shayok Dutta (Rice University), Antonio Fernandez-Ruiz (Cornell University) and Josh Siegle (Allen Institute) for sharing data used in example analyses. The development of GhostiPy was supported by the National Science Foundation (NSF CBET1351692) and the National Institute of Neurological Diseases and Strokes (R01NS115233).

## References

[1] Hemant Bokil et al. “Chronux: a platform for analyzing neural signals”. In: Journal of neuroscience methods 192.1 (2010), pp. 146–151.

[2] C Sidney Burrus, Admadji W Soewito, and Ramesh A Gopinath. “Least squared error FIR filter design with transition bands”. In: IEEE Transactions on Signal Processing 40.6 (1992), pp. 1327–1340.

[3] Ryan T Canolty et al. “High gamma power is phase-locked to theta oscillations in human neocortex”. In: science 313.5793 (2006), pp. 1626–1628.

[4] Ingrid Daubechies. “A nonlinear squeezing of the continuous wavelet transform based on auditory nerve models”. In: Wavelets in medicine and biology (1996), pp. 527–546.

[5] Ingrid Daubechies, Yi Wang, and Hau-tieng Wu. “ConceFT: Concentration of frequency and time via a multitapered synchrosqueezed transform”. In: Philosophical Transactions of the Royal Society A: Mathematical, Physical and Engineering Sciences 374.2065 (2016), p. 20150193.

[6] Intel developers. Performance Benchmarks. https://software.intel.com/content/www/us/en/develop/tools/math-kernel-library/benchmarks.html. [Online; accessed 2020-06-09]. 2020.

[7] Dino Dvorak and André A Fenton. “Toward a proper estimation of phase–amplitude coupling in neural oscillations”. In: Journal of Neuroscience methods 225 (2014), pp. 42–56.

[8] Kelly R Fitz and Sean A Fulop. “A unified theory of time-frequency reassignment”. In: arXiv preprint arXiv:0903.3080 (2009).

[9] Matteo Frigo and Steven G Johnson. “FFTW: An adaptive software architecture for the FFT”. In: Proceedings of the 1998 IEEE International Conference on Acoustics, Speech and Signal Processing, ICASSP’98 (Cat. No. 98CH36181). Vol. 3. IEEE. 1998, pp. 1381–1384.

[10] Matteo Frigo and Steven G Johnson. “The design and implementation of FFTW3”. In: Proceedings of the IEEE 93.2 (2005), pp. 216–231.

[11] Timothy J Gardner and Marcelo O Magnasco. “Sparse time-frequency representations”. In: Proceedings of the National Academy of Sciences 103.16 (2006), pp. 6094–6099.

[12] Alexandre Gramfort et al. “MEG and EEG data analysis with MNE-Python”. In: Frontiers in neuroscience 7 (2013), p. 267.

[13] Gregory Lee et al. “PyWavelets: A Python package for wavelet analysis”. In: Journal of Open Source Software 4.36 (2019), p. 1237.

[14] JM Lilly and J-C Gascard. “Wavelet ridge diagnosis of time-varying elliptical signals with application to an oceanic eddy”. In: Nonlinear Processes in Geophysics 13.5 (2006), pp. 467–483.

[15] Jonathan M Lilly and Sofia C Olhede. “Generalized Morse wavelets as a superfamily of analytic wavelets”. In: IEEE Transactions on Signal Processing 60.11 (2012), pp. 6036–6041.

[16] Jonathan M Lilly and Sofia C Olhede. “Higher-order properties of analytic wavelets”. In: IEEE Transactions on Signal Processing 57.1 (2008), pp. 146–160.

[17] Sofia C Olhede and Andrew T Walden. “Generalized morse wavelets”. In: IEEE Transactions on Signal Processing 50.11 (2002), pp. 2661–2670.

[18] Robert Oostenveld et al. “FieldTrip: open source software for advanced analysis of MEG, EEG, and invasive electrophysiological data”. In: Computational intelligence and neuroscience 2011 (2011).

[19] Donald B Percival, Andrew T Walden, et al. Spectral analysis for physical applications. cambridge university press, 1993.

[20] M. Reinecke. Performance Benchmarks. https://gitlab.mpcdf.mpg.de/mtr/pocketfft. [Online; accessed 2020-06-11]. 2019.

[21] Matthew Rocklin. “Dask: Parallel computation with blocked algorithms and task scheduling”. In: Proceedings of the 14th python in science conference. Vol. 126. Citeseer. 2015.

[22] Franśois Tadel et al. “MEG/EEG group analysis with brainstorm”. In: Frontiers in neuroscience 13 (2019), p. 76.

[23] Jeffery L Teeters et al. “Neurodata without borders: creating a common data format for neurophysiology”. In: Neuron 88.4 (2015), pp. 629–634.

[24] Gaurav Thakur et al. “The synchrosqueezing algorithm for time-varying spectral analysis: Robustness properties and new paleoclimate applications”. In: Signal Processing 93.5 (2013), pp. 1079–1094.

[25] David J Thomson. “Spectrum estimation and harmonic analysis”. In: Proceedings of the IEEE 70.9 (1982), pp. 1055–1096.

[26] Stefan Van Der Walt, S Chris Colbert, and Gael Varoquaux. “The NumPy array: a structure for efficient numerical computation”. In: Computing in science & engineering 13.2 (2011), pp. 22–30.

[27] Pauli Virtanen et al. “SciPy 1.0: fundamental algorithms for scientific computing in Python”. In: Nature methods 17.3 (2020), pp. 261–272.

[28] Alper Yegenoglu et al. Elephant–open-source tool for the analysis of electrophysiological data sets. Tech. rep. Computational and Systems Neuroscience, 2015.

