## Supplementary material for "GhostiPy: an efficient signal processing and spectral analysis toolbox for large data": Source code: analytic.py

```
from multiprocessing import cpu_count
import numpy as np
import pyfftw
import gc

def _check_params(signal, axis, pad_to, threads):
    if not np.all(np.isreal(signal)):
        raise ValueError("The input data must be real-valued")

    if np.any(np.array(signal.shape) == 0): 
        raise ValueError("Cannot do computation on an empty array")

    if np.abs(axis) > signal.ndim:
        raise ValueError(f"Specified axis {axis} is out-of-bounds for input"
                         f" signal with {signal.ndim} dimensions")

    if threads < 1:
        raise ValueError("Must have at least one thread to do the computation...")

    L = signal.shape[axis]
    
    if pad_to is not None:
        if pad_to < L:
            raise ValueError("'pad_to' must be at least the length of the"
                             " input data")

def analytic_signal(signal, *, axis=-1, pad_to=None, threads=cpu_count()):

    """Computes the analytic signal x_a = x + iy where
    x is the original signal, and y is the Hilbert transform
    of x

    Parameters
    ----------
    signal : numpy.ndarray
        Must be 1D and real
    axis : int, optional
        Axis along which to compute the analytic signal
    pad_to : float, optional
        Zero-pad signal along the specified axis to this length.
        Note that at the end, the returned array is truncated
        to remove the padding once the analytic signal is computed.
        The output is thus the same shape as the input.
        Generally it is advised to avoid padding.
    threads : int, optional
        How many threads to use.
        Default is the total number of virtual threads, which may
        be larger than the number of physical CPU cores.

    Returns
    -------
    A 1D array containing the analytic signal
    """
    
    if pad_to is None:
        pad_to = signal.shape[axis]
    
    _check_params(signal, axis, pad_to, threads)
        
    nfft = pad_to
    L = signal.shape[axis]
    M = nfft // 2 + 1

    # Generating the analytic signal involves taking the FT of the signal
    # and zeroing out the negative frequencies. Therefore, we don't
    # actually need to compute the negative frequencies. Instead, we
    # do a real-valued FFT to get the positive frequencies only. Note that
    # the signal must be real to use this strategy correctly!
    xtd_dims = [val for val in signal.shape]
    xfd_dims = [val for val in signal.shape]
    xtd_dims[axis] = nfft
    xfd_dims[axis] = M

    xtd = pyfftw.zeros_aligned(tuple(xtd_dims), dtype='float64')
    xfd = pyfftw.zeros_aligned(tuple(xfd_dims), dtype='complex128')
    fft_sig = pyfftw.FFTW( 
        xtd, xfd,
        axes=(axis, ),
        direction='FFTW_FORWARD',
        flags=['FFTW_ESTIMATE'],
        threads=threads,
        planning_timelimit=0 )
    
    # We don't care about the spectrum of the analytic signal
    # so we can compute an in-place FFT to save memory
    a = pyfftw.zeros_aligned(tuple(xtd_dims), dtype='complex128')
    fft_analytic_inv = pyfftw.FFTW(
        a, a,
        axes=(axis, ),
        direction='FFTW_BACKWARD',
        flags=['FFTW_ESTIMATE'], 
        threads=threads,
        planning_timelimit=0 )

    xtd_slices = [np.s_[:]] * signal.ndim
    xtd_slices[axis] = np.s_[0:L]
    xtd[tuple(xtd_slices)] = signal
    fft_sig(normalise_idft=True)

    a_slices = [np.s_[:]] * a.ndim
    xfd_slices = [np.s_[:]] * xfd.ndim

    a_slices[axis] = np.s_[0:1]
    # This is true regardless if the DFT is odd or even length
    a[tuple(a_slices)] = xfd[tuple(a_slices)]

    if nfft & 1:  # odd
        a_slices[axis] = np.s_[1:(nfft + 1)//2]
        xfd_slices[axis] = np.s_[1:]

        a[tuple(a_slices)] = 2*xfd[tuple(xfd_slices)]
    else:
        # The last value in an even-length real-valued DFT is both
        # positive and negative so we make sure to keep it in addition
        # to DC
        a_slices[axis] = np.s_[nfft//2:nfft//2 + 1]
        xfd_slices[axis] = np.s_[-1:]
        a[tuple(a_slices)] = xfd[tuple(xfd_slices)]
        
        a_slices[axis] = np.s_[1:nfft//2]
        xfd_slices[axis] = np.s_[1:-1]
        a[tuple(a_slices)] = 2*xfd[tuple(xfd_slices)]

    # No longer used, so remove now to save memory
    del xtd
    del xfd
    gc.collect()

    fft_analytic_inv(normalise_idft=True)

    return a[tuple(xtd_slices)]

def signal_envelope(signal, *, axis=-1, pad_to=None, threads=cpu_count()):
    """Computes the envelope of a signal by determining the analytic signal
    x_a = x + iy (where x is the original signal, and y is the Hilbert transform
    of x), and taking the magnitude of x_a at every point.

    Parameters
    ----------
    signal : numpy.ndarray
        Must be 1D and real
    axis : int, optional
        Axis along which to compute the analytic signal
    pad_to : float, optional
        Zero-pad signal along the specified axis to this length.
        Note that at the end, the returned array is truncated
        to remove the padding once the analytic signal is computed.
        The output is thus the same shape as the input.
        Generally it is advised to avoid padding.
    threads : int, optional
        How many threads to use.
        Default is the total number of virtual threads, which may
        be larger than the number of physical CPU cores.

    Returns
    -------
    A 1D array containing the analytic signal
    """

    _check_params(signal, axis, pad_to, threads)
    
    return np.abs(
        analytic_signal(
            signal,
            axis=axis,
            pad_to=pad_to,
            threads=threads)
        )
    
def signal_phase(signal, *, axis=-1, pad_to=None, threads=cpu_count()):
    """Computes the instantaneous phase of a signal.

    Parameters
    ----------
    signal : numpy.ndarray
        Must be 1D and real
    axis : int, optional
        Axis along which to compute the analytic signal
    pad_to : float, optional
        Zero-pad signal along the specified axis to this length.
        Note that at the end, the returned array is truncated
        to remove the padding once the analytic signal is computed.
        The output is thus the same shape as the input.
        Generally it is advised to avoid padding.
    threads : int, optional
        How many threads to use.
        Default is the total number of virtual threads, which may
        be larger than the number of physical CPU cores.

    Returns
    -------
    A 1D array containing the analytic signal
    """

    _check_params(signal, axis, pad_to, threads)

    return np.angle(
        analytic_signal(
            signal,
            axis=axis,
            pad_to=pad_to,
            threads=threads)
        )
```
