## Supplementary material for "GhostiPy: an efficient signal processing and spectral analysis toolbox for large data": Source code: convolution.py

```
import numpy as np
import pyfftw
from multiprocessing import cpu_count
import time

def osconvolve(signal, kernel, *, mode='full', nfft=None,
               threads=cpu_count(), axis=-1, outarray=None,
               input_index_bounds=None, output_index_bounds=None,
               describe_dims=False, ds=None, input_dim_restrictions=None,
               output_offset=0, verbose=False):
    """Computes a convolution with FFTW, using the convolution
    theorem. This function has been written to minimize memory
    usage.
    """
    if np.isrealobj(signal) and np.isrealobj(kernel):
        real_output = True
        
    if threads < 1:
        raise ValueError("Must have at least one thread to do the FFT...")
        
    if kernel.ndim != 1:
        raise ValueError("Kernel must be 1D")
    
    ###############################################################
    # Determine convolution lengths
    if input_index_bounds is not None:
        if len(input_index_bounds) != 2:
            raise ValueError(f"Got {len(input_index_bounds)} elements in"
                             " input_index_bounds but must have 2 elements")
        if not input_index_bounds[0] < input_index_bounds[1]:
            raise ValueError(f"'input_index_bounds' {input_index_bounds} is not"
                             f" strictly increasing")
        try:
            test_slices = [np.s_[:]] * signal.ndim
            test_slices[axis] = input_index_bounds[0]
            test_1 = signal[tuple(test_slices)]
            test_slices[axis] = input_index_bounds[1]
            test_2 = signal[tuple(test_slices)]
        except:
            raise IndexError(f"input_index_bounds are out of bounds for"
                             " input array")
        N = input_index_bounds[1] - input_index_bounds[0]
    else:
        N = signal.shape[axis]
    
    M = kernel.shape[0]
    tot_length = N + M - 1
    
    if mode == 'full':
        outsize = tot_length
    elif mode == 'same':
        outsize = N
    elif mode == 'valid':
        if N < M:
            raise ValueError("Cannot do a 'valid' convolution because "
                             "the input is shorter than the kernel")
        outsize = N - M + 1
    else:
        raise ValueError(f"Got invalid value {mode} for 'mode'")
    
    if output_index_bounds is not None:
        if mode != 'full':
            raise NotImplementedError("'output_index bounds' currently implemented only"
                                      " for mode == 'full'")
        if len(output_index_bounds) != 2:
            raise ValueError(f"Got {len(output_index_bounds)} elements in"
                             " 'output_index_bounds' but must have 2 elements")
        if (output_index_bounds[0] < 0 or output_index_bounds[0] >= tot_length or
            output_index_bounds[1] < 0 or output_index_bounds[1] >= tot_length):
            raise ValueError(f"'output_index_bounds' can only be in the range"
                             f" [{0}, {tot_length}], inclusive")
        if not output_index_bounds[0] < output_index_bounds[1]:
            raise ValueError(f"'output_index_bounds' {output_index_bounds} is not"
                             f" strictly increasing")

        first_ind = output_index_bounds[0]
        last_ind = output_index_bounds[1]
    else:
        first_ind = (tot_length - outsize) // 2
        last_ind = first_ind + outsize
#     print(f"First ind {first_ind}, last ind {last_ind}, diff {last_ind - first_ind}, total size {tot_length}")
    
    downsample = False
    if ds is not None:
        if divmod(ds, 1)[1] != 0:
            raise ValueError("'ds' factor must be an integer > 1")
        ds = int(ds)
        block_offset = 0
        downsample = True
    
    ###############################################################
    # handle output params
    if real_output:
        dtype = '<f8'
    else:
        dtype = '<c16'
    
    # set the default expected_shapes...
    expected_shape = [dimsize for dimsize in signal.shape]
    expected_shape[axis] = last_ind - first_ind
    
    # ... and override the appropriate part if input_dim_restrictions
    # were passed in
    if input_dim_restrictions is not None:
        if len(input_dim_restrictions) != signal.ndim:
            raise ValueError(f"Expected {signal.ndim} elements in 'input_dim_restrictions'"
                             f" but got {len(input_dim_restrictions)}")
        if input_dim_restrictions[axis] is not None:
            raise ValueError(f"input_dim_restrictions[{axis}] must be set to None")
        for dim in range(signal.ndim):
            temp_slice = input_dim_restrictions[dim]
            if temp_slice is not None:
                expected_shape[dim] = len(temp_slice)
    
    # continue modifying expected_shape if downsampling
    if downsample:
        n, mod = divmod(last_ind - first_ind, ds)
        if mod != 0:
            n += 1
        expected_shape[axis] = n
    
    expected_shape = tuple(expected_shape)
    if describe_dims:
        if verbose:
            print(f"Output array should have shape {expected_shape} and dtype {dtype}")
        return expected_shape, dtype

    if outarray is None:
        outarray = np.zeros(expected_shape, dtype=dtype)
        if verbose:
            print(f"Allocated array of shape {outarray.shape} with dtype {dtype}")
    else:
        if verbose:
            print(
                "Checking output array shape is disabled, make sure portion of"
                f" output array has shape {expected_shape}")
        if not real_output and np.isrealobj(outarray[..., 0:1]):
            raise TypeError("Output array is real but expected one of a complex dtype")

    ######################################################################
    if nfft is not None:
        if nfft < M:
            raise ValueError(f"'nfft' must be at least the kernel"
                             f" size of {M}")
    else:   # Choose good default fft_length
        nfft = 65536
        while nfft < 10 * M:
            nfft *= 4
            
    L = nfft - (M - 1)

    #############################################################################
    x_dims = [dimsize for dimsize in signal.shape]
    x_dims[axis] = nfft
    if input_dim_restrictions is not None:
        for dim in range(signal.ndim):
            temp_slice = input_dim_restrictions[dim]
            if temp_slice is not None:
                x_dims[dim] = len(temp_slice)
    
    
    # Use in-place transform
    x = pyfftw.zeros_aligned(tuple(x_dims), dtype='complex128')
    fft_sig = pyfftw.FFTW( x, x,
                           axes=(axis, ),
                           direction='FFTW_FORWARD',
                           flags=['FFTW_ESTIMATE'], 
                           threads=threads)

    
    ##############################################################################
    y_dims = [1] * signal.ndim
    y_dims[axis] = nfft
    
    y = pyfftw.zeros_aligned(tuple(y_dims), dtype='complex128')
    fft_kernel = pyfftw.FFTW( y, y,
                              axes=(axis, ),
                              direction='FFTW_FORWARD', 
                              flags=['FFTW_ESTIMATE'], 
                              threads=threads)

    y_slices = [0] * signal.ndim
    y_slices[axis] = np.s_[0:M]
    y[tuple(y_slices)] = kernel
    y_slices[axis] = np.s_[M:nfft]
    y[tuple(y_slices)] = 0
    # Notice that once we take the FFT of the kernel, we never have to
    # do it again! Just multiply with the FFT of each chunk. This saves
    # us computation
    fft_kernel()
#     assert np.allclose(y, fft(kernel, fft_length))
    ################################################################################

    # We don't care about the convolution looks like in the frequency
    # domain so we can compute an in-place FFT to save memory (and
    # potentially computation time)
    conv_dims = [dimsize for dimsize in signal.shape]
    conv_dims[axis] = nfft
    if input_dim_restrictions is not None:
        for dim in range(signal.ndim):
            temp_slice = input_dim_restrictions[dim]
            if temp_slice is not None:
                conv_dims[dim] = len(temp_slice)
    conv = pyfftw.zeros_aligned(tuple(conv_dims), dtype='complex128')
    fft_conv_inv = pyfftw.FFTW(conv, conv,
                               axes=(axis, ),
                               direction='FFTW_BACKWARD', 
                               flags=['FFTW_ESTIMATE'], 
                               threads=threads)
    
    ####################################################################
    
    start_offset = 0
    if input_index_bounds is not None:
        start_offset = input_index_bounds[0]
    div, _ = divmod(tot_length, L)
    bounds = np.arange(start_offset, start_offset + (div+2)*L, L)
    block_bounds = np.vstack((bounds[:-1], bounds[1:])).T
    n_blocks = block_bounds.shape[0]
    
    signal_slices_1 = [np.s_[:]] * signal.ndim
    signal_slices_2 = [np.s_[:]] * signal.ndim
    if input_dim_restrictions is not None:
        for dim in range(signal.ndim):
            temp_slice = input_dim_restrictions[dim]
            if temp_slice is not None:
                signal_slices_1[dim] = temp_slice
                signal_slices_2[dim] = temp_slice
    
    x_slices_1 = [np.s_[:]] * x.ndim
    x_slices_2 = [np.s_[:]] * x.ndim
    
    outarray_slices = [np.s_[:]] * outarray.ndim
    conv_slices = [np.s_[:]] * conv.ndim
    
    outarray_marker = output_offset
    
    # note that first_ind and last_ind are used to index into
    # the output of the convolution. They are not used to determine
    # where in this function's output array the convolution results
    # are written. The outarray_marker keeps track of that.
    first_block_to_check, first_offset = divmod(first_ind, L)
    last_block_to_check, last_offset = divmod(last_ind, L)
    # print(f'first index {first_ind}, last index {last_ind}')
    # print(f"First segment: {M-1}, second segment {L}, for a total of {nfft} FFT points")
    # print(f"Total number of blocks: {n_blocks}")
    # print(f'Index {first_ind}, first block to check: {first_block_to_check}')
    # print(f"Index {last_ind} last block to check: {last_block_to_check}")
    # print(f'First offset {first_offset}, last offset {last_offset}')

    tot_samples = 0
    tr = 0 # read time
    tp = 0 # processing time
    tw = 0 # write time

    for ii, (start, stop) in enumerate(block_bounds):
        t0 = time.time()

        if ii < first_block_to_check or ii > last_block_to_check:
            pass
        else:
            tr0 = time.time()
            # print(f"Computing block {ii} of {n_blocks - 1}")
            # initialize entire block to 0, then fill with
            # appropriate input data if it exists
            x[:] = 0 
            ind1 = start - (M-1)
            # fill M - 1 portion of block
            # M - 1 segment extends past start of data,
            # but may overlap partially with the data.
            # Make sure to put this part at the END
            # of the M - 1 segment
            if ind1 < 0:
                ind1 = 0
                signal_slices_1[axis] = np.s_[ind1:start]
                signal_chunk = signal[tuple(signal_slices_1)]
                length = signal_chunk.shape[axis]
                diff = M - 1 - length
                x_slices_1[axis] = np.s_[diff:M-1]
                x[tuple(x_slices_1)] = signal_chunk
            # Start of M - 1 segment has valid data, though not
            # guaranteed to fill entire portion if extends beyond
            # data end boundary
            else:
                try:
                    signal_slices_1[axis] = np.s_[ind1:start]
                    signal_chunk = signal[tuple(signal_slices_1)]
                    length = signal_chunk.shape[axis]
                    x_slices_1[axis] = np.s_[:length]
                    x[tuple(x_slices_1)] = signal_chunk
                except:
                    # M - 1 segment already initialized to 0, so leave as is
#                     print(f"Requested {ind1} out of bounds, doing nothing")
                    pass

            # fill L segment of block
            signal_slices_2[axis] = np.s_[start:stop]
            signal_chunk = signal[tuple(signal_slices_2)]
            length = signal_chunk.shape[axis]
            x_slices_2[axis] = np.s_[M-1:M-1 + length]

            x[tuple(x_slices_2)] = signal_chunk
            tr += time.time() - tr0

            tp0 = time.time()
            fft_sig(normalise_idft=True)
            conv[:] = x * y
            fft_conv_inv(normalise_idft=True)

            if ii == first_block_to_check and ii == last_block_to_check:

                if downsample:
                    conv_slices[axis] = np.s_[M-1+first_offset:M-1+last_offset:ds]
                    n_samples = conv[tuple(conv_slices)].shape[axis]
                    outarray_slices[axis] = np.s_[outarray_marker:outarray_marker+n_samples]
                else:
                    n_samples = last_offset - first_offset
                    conv_slices[axis] = np.s_[M-1+first_offset:M-1+last_offset]
                    outarray_slices[axis] = np.s_[outarray_marker:outarray_marker+n_samples]

            elif ii == first_block_to_check:

                if downsample:
                    conv_slices[axis] = np.s_[M-1+first_offset::ds]
                    
                    max_samples = L - first_offset
                    n_samples, rem = divmod(max_samples, ds)
                    if rem != 0:
                        n_samples += 1
                        block_offset = ds - rem
                    else:
                        block_offset = 0
#                     print(f"Wrote {n_samples} samples. Leftover {rem}. Next offset {block_offset}")
                    outarray_slices[axis] = np.s_[outarray_marker:outarray_marker+n_samples]
                else:
                    n_samples = nfft - (M - 1 + first_offset)
#                     print(f'Block {ii}, taking {n_samples} samples')
                    conv_slices[axis] = np.s_[M - 1 + first_offset:nfft]
                    outarray_slices[axis] = np.s_[outarray_marker:outarray_marker+n_samples]

            elif ii == last_block_to_check:

                if downsample:
#                     print("last block to check!")
#                     print(f"Searching range {M-1+block_offset} to {M-1+last_offset}")
                    conv_slices[axis] = np.s_[M-1+block_offset:M-1+last_offset:ds]
                    n_samples = conv[tuple(conv_slices)].shape[axis]
                    outarray_slices[axis] = np.s_[outarray_marker:outarray_marker+n_samples]
                else:
#                     print(f"Block {ii}, last block to check")
                    n_samples = last_offset
#                     print(f'Block {ii}, taking {n_samples} samples')
#                     n_samples = outarray.shape[axis] - outarray_marker
#                     print(f"Searching range {M-1} to {M-1+n_samples}")
                    conv_slices[axis] = np.s_[M-1:M-1 + n_samples]
                    outarray_slices[axis] = np.s_[outarray_marker:outarray_marker + n_samples]

            else:

                if downsample:
                    conv_slices[axis] = np.s_[M-1+block_offset:nfft:ds]
                    
                    max_samples = L - block_offset
                    n_samples, rem = divmod(max_samples, ds)
                    if rem != 0:
                        n_samples += 1
                        block_offset = ds - rem
                    else:
                        block_offset = 0
                    outarray_slices[axis] = np.s_[outarray_marker:outarray_marker + n_samples]
#                     print(f"Wrote {n_samples} samples. Leftover {rem}. Next offset {block_offset}")
                else:
#                     print(f'Block {ii} of {n_blocks-1}, taking {n_samples} samples')
                    n_samples = stop - start
                    outarray_slices[axis] = np.s_[outarray_marker:outarray_marker + n_samples]
                    conv_slices[axis] = np.s_[M-1:nfft]

            tp += time.time() - tp0

            tw0 = time.time()
            if real_output:
                outarray[tuple(outarray_slices)] = conv[tuple(conv_slices)].real
            else:
                outarray[tuple(outarray_slices)] = conv[tuple(conv_slices)]
            outarray_marker += n_samples
            tot_samples += n_samples
            tw += time.time() - tw0

        if verbose:
            print(
                f"Computed block {ii} of {n_blocks - 1}, "
                f"elapsed time: {time.time() - t0} seconds")

#     print(f"Wrote {tot_samples} data points")
    if not expected_shape[axis] == tot_samples:
        raise ValueError(f"Expected to write {expected_shape[axis]} samples "
                         f"for axis {axis} but actually wrote {tot_samples}")

    if verbose:
        print(f"Total read time: {tr} seconds")
        print(f"Total processing time: {tp} seconds")
        print(f"Total write time: {tw} seconds")
    
    return outarray
```
