## Supplementary material for "GhostiPy: an efficient signal processing and spectral analysis toolbox for large data": Source code: filtering.py

```
from ghostipy.dsp.convolution import osconvolve
from multiprocessing import cpu_count

__all__ = ['filter_data_fir']

def filter_data_fir(data, b, *, nfft=None,
                    threads=cpu_count(), axis=-1, outarray=None,
                    input_index_bounds=None, output_index_bounds=None,
                    describe_dims=False, ds=None, input_dim_restrictions=None,
                    output_offset=0, verbose=False):
    """
    Applies an FIR filter to data.

    Parameters
    ----------
    data : array-like
        The data to be filtered
    b : array-like
        Filter coefficients
    nfft : integer, optional
        Number of elements to use for the FFT, for the axis along which
        the filter is applied.
        Default is chosen by convolution routine used to filter the data.
    threads : integer, optional
        Number of threads to use for the FFT.
        Default is the total number of CPU cores (which may be virtual).
    axis : integer, optional
        Axis along which to apply the filter.
        Default is -1, the last axis.
    outarray : numpy.ndarray, optional
        The output array.
        Default is None, in which case the output array is created while
        executing this method.
    input_index_bounds : array-like with 2 elements, optional
        An array containing two elements, denoting the start (inclusive)
        and stop (exclusive) indices of the input data for the specified
        axis. 
        Default is None, in which case the entire input data for the
        specified axis are used.
    output_index_bounds : array-like with 2 elements, optional
        An array containing two elements, denoting the start (inclusive)
        and stop (exclusive) indices of the output data for the specified
        axis. 
        Default is None, in which case this method returns the full output
        from applying the filter.
    describe_dims : boolean, optional
        Whether to return the expected shape and dtype of the output and
        return immediately (no filtering is performed). This option is
        useful for out-of-core computation. While the expected shape
        should not be changed, the dtype is only suggested, e.g. it is
        acceptable to use a lower precision dtype (such as float32 instead
        of float64 to save space)
        Default is False.
    ds : integer, optional
        Downsampling factor.
        Default is None, in which case the output is not downsampled
    input_dim_restrictions : array-like or None, optional
        Array with number of elements equal to the number of dimensions
        of the input data. Each element in 'input_slice_params' can either
        be None, or an array-like specifying which indices to use, under
        the requirement that input_slice_params[axis] must be None.
        Default is None, in which case all the input data are used.
    output_offset : integer, optional
        Index at which to begin storing the output of this method.
        This offset is only applied to the axis along which the data
        are filtered.
        Default is 0.
    verbose : bool, optional
        Whether to show more output while filtering.
        Default is False.

    Returns
    -------
    If 'describe_dims' is True:
        shape, dtype : tuple
            Expected output array shape and dtype
    Otherwise:
        outarray : array-like
            The filtered data  


    """

    return osconvolve(
        data,
        b,
        mode='full',
        nfft=nfft,
        threads=threads,
        axis=axis,
        outarray=outarray,
        input_index_bounds=input_index_bounds,
        output_index_bounds=output_index_bounds,
        describe_dims=describe_dims,
        ds=ds,
        input_dim_restrictions=input_dim_restrictions,
        output_offset=output_offset,
        verbose=verbose)
```
