## Supplementary material for "GhostiPy: an efficient signal processing and spectral analysis toolbox for large data": Source code: firfilter.py

```
import numpy as np

__all__ = ['estimate_taps',
           'firdesign',
           'group_delay']

def group_delay(b):
    """
    Gives the group delay of a linear phase FIR Type I filter. Useful for
    ghostipy.filter_data_fir()

    Parameters
    ----------
    b : np.ndarray, shape (N,)
        The filter coefficients

    Return
    ------
    K : int
        The group delay
    """

    L = len(b)
    if not L & 1:
        raise ValueError(
            "There are {} filter coefficients (an even number), so the group delay "
            "cannot be converted to an integer value".format(L))
    
    return (L - 1) // 2

def estimate_taps(fs, tw, *, d1=None, d2=None):
    
    """
    Estimates number of taps needed to achieve desired frequency response of
    a Type I FIR filter

    Parameters
    ----------
    fs : scalar
        Sampling rate in Hz
    tw : scalar
        Transition bandwidth in Hz
    d1 : scalar
        Passband deviation, optional
        Default is 0.1% (0.001)
    d2 : scalar
        Minimum stopband attenuation, optional
        Default is 60 dB (1e-6)
    
    Returns
    -------
    numtaps : integer
        Number of taps. Will be an odd number.
    
    References
    ----------
    https://dsp.stackexchange.com/questions/31066/how-many-taps-does-an-fir-filter-need/31077
    """
    
    if d1 is None:
        d1 = 1e-3
    if d2 is None:
        d2 = 1e-6

    numtaps = int(np.ceil(2/3*np.log10(1/(10*d1*d2))*fs/(tw)))
    if not numtaps & 1:
        numtaps += 1
    return numtaps

def _firspline(numtaps, f1, f2, *, fs=None, p=None):
    
    """
    Designs a Type I low-pass filter with splines in the transition band

    Parameters
    ----------
    numtaps : number of coefficients
    f1 : Frequency where the filter's amplitude response is 1, in Hz
    f2 : Frequency where the filter's amplitude response is 0, in Hz
    p : Spline power, optional.
        Default is 0.312*numtaps*(f2 - f1)/nyq. Empirical formula, see
        Burrus et al, 1992
    fs : Sampling frequency in Hz, optional.
        Default is 2 Hz
    
    Returns
    -------
    b : The filter coefficients
    """
    
    if not numtaps & 1:
        raise ValueError("numtaps must be odd but got {}".format(numtaps))

    if fs is None:
        fs = 2
    nyq = fs/2

    if f1 > nyq or f2 > nyq:
        raise ValueError("Got critical frequencies {} and {} but they must both"
                         " be less than the Nyquist frequency of {}".
                         format(f1, f2, nyq))

    if not f1 < f2:
        raise ValueError("f1 must be <{} but got {}".format(f2, f1))

    if p is None:
        p = 0.312*numtaps*(f2 - f1)/nyq
        # print(p)
    if not p >= 0:
        raise ValueError("p must be positive but got {}".format(p))
        
    # Convert to normalized frequency in radians
    wo = (f1 + f2)/2 * (1/nyq * np.pi)
    dw = (f2 - f1)/2 * (1/nyq * np.pi)
#     print(wo, dw)
#     print("Spline power is {}".format(p))
 
#     if fs is None:
#         fs = 2
#     p1 = 0.62*(f2-f1)*(fs/2)*N
#     print("Spline power from Hz is {}".format(p1))

    nvec = np.arange(1, (numtaps-1)/2 + 1)
    
    h = np.sin(wo*nvec) / (np.pi*nvec) # Optimal L2 solution to ideal lowpass filter

    x = dw * nvec / p
    spline = (np.sin(x) / x) ** p
    h *= spline  # connect transition band with spline
    
    b = np.hstack((np.flip(h), wo/np.pi, h))  # make linear phase
    
    return b

def firdesign(numtaps, band_edges, desired, *, fs=1, p=None):
    """
    Designs an arbitrary Type I FIR filter with spline transition
    bands, optimized for an L2 error norm

    Parameters
    ----------
    numtaps : scalar, integer
        Number of filter coefficients. Must be an odd number.
    band_edges : array-like
        Critical frequencies of filter. Do not include 0 or Nyquist.
    desired : array-like
        Magnitude response of filter at specified band_edges. Can only
        be 0 or 1.
    fs : scalar, optional
        Sampling rate of filter, in Hz.
        Default is 1 Hz.
    p : scalar, optional
        Power to which the spline transition band functions are raised.
        Default is None, in which case it is determined according to
        Burrus et al, 1992.

    Returns
    -------
    b : numpy.ndarray
        The filter coefficients
    """
    
    band_edges = np.array(band_edges)
    desired = np.array(desired)

    if not numtaps & 1:
        raise ValueError(f"Got {numtaps} for 'numtaps' but must be an odd value")
    if not len(band_edges) % 2 == 0:
        raise ValueError("Must have even number of band edges")
    if not len(band_edges) == len(desired):
        raise ValueError("must have equal number of band edges and values")
    if not np.sum((desired == 0) + (desired == 1)) == len(desired):
        raise ValueError("All values must be either 0 or 1")
    if not band_edges[0] > 0:
        raise ValueError("First band edge must be greater than 0")
    if not band_edges[-1] < fs / 2:
        raise ValueError(f"Last band edge must be less than {fs/2}")
    if not np.all(band_edges[:-1] < band_edges[1:]):
        raise ValueError("'band_edges' must be a monotonically increasing sequence")

    desired_pairs = np.vstack((desired[:-1], desired[1:])).T
    band_pairs =  np.vstack((band_edges[:-1], band_edges[1:])).T
    for pair_ind, ((edge1, edge2), (v1, v2)) in enumerate(zip(band_pairs, desired_pairs)):
        if pair_ind % 2 == 0:
            if v1 == v2:
                raise ValueError(f"Got {v1} for band edge {edge1} Hz"
                                 f" and {v2} for band edge {edge2} Hz but"
                                 f" must be different values")
        else:
            if v1 != v2:
                raise ValueError(f"Got {v1} for band edge {edge1} Hz"
                                 f"and {v2} for band edge {edge2} Hz but"
                                 " must be the same values")

    critical_points = band_edges.reshape((-1, 2))
    # low pass prototypes
    prototypes = np.zeros((critical_points.shape[0], numtaps))
    for ind, (f1, f2) in enumerate(critical_points):
        prototypes[ind] = _firspline(numtaps, f1, f2, fs=fs, p=p)

    if prototypes.shape[0] == 1: # single band
        b = prototypes[0]
        if desired[-1] == 1: # high pass
            cr = np.hstack((np.zeros((numtaps-1)//2), 1, np.zeros((numtaps-1)//2)))
            b = cr - prototypes[0]

    else: # multi-band
        b = np.zeros(numtaps)
        
        # Magnitude at 0 and Nyquist is the same
        if desired[0] == desired[-1]:
            for ii in np.arange(prototypes.shape[0], step=2):
                bl = prototypes[ii]
                bh = prototypes[ii+1]
                b += (bh - bl)

            # high pass at 0 and Nyquist, so invert
            if desired[-1] == 1:
                cr = np.hstack((np.zeros((numtaps-1)//2), 1, np.zeros((numtaps-1)//2)))
                b = cr - b


        else:
            if desired[0] == 0:
                tmp = prototypes[-1]
                cr = np.hstack((np.zeros((numtaps-1)//2), 1, np.zeros((numtaps-1)//2)))
                b_special = cr - tmp
                prototypes = prototypes[:-1]
            else:
                b_special = prototypes[0]
                prototypes = prototypes[1:]

            for ii in np.arange(prototypes.shape[0] - 1, step=2):
                bl = prototypes[ii]
                bh = prototypes[ii+1]
                b += (bh - bl)
            
            b += b_special

    return b
```
