## Supplementary material for "GhostiPy: an efficient signal processing and spectral analysis toolbox for large data": Source code: core.py

```
import numpy as np
import xarray as xr
import holoviews as hv
import hvplot.xarray
import dask.array as da
import matplotlib.pyplot as plt
from multiprocessing import cpu_count

__all__ = ['plot_fourier_spectrogram',
           'plot_wavelet_spectrogram',
           'plot_frequency_response']
        
def plot_fourier_spectrogram(s, f, t, *, freq_limits=None, time_limits=None,
                             zscore=False, zscore_axis=None, n_workers=cpu_count(),
                             timescale='seconds', relative_time=False, center_time=False,
                             colorbar=True, plot_type='pcolormesh', fig=None,
                             ax=None, **kwargs):
    """
    Plot a Fourier based spectrogram. Note that the spectrogram
    data need not fit into memory but the data used for the
    plot should.

    Parameters
    ----------
    s : np.ndarray, with shape (n_freqs, n_timepoints)
        The spectrogram. Should be real-valued.
    f : np.ndarray, with shape (n_freqs, )
        Spectrogram frequencies in Hz. Must be uniformly
        spaced and strictly increasing or strictly
        decreasing.
    t : np.ndarray, with shape (n_timepoints, )
        Midpoints of each spectrogram time window in seconds.
        Must be uniformly spaced and strictly increasing
        or strictly decreasing.
    freq_limits : list or None, optional
        If not None, a list consisting of the half-open interval
        [min_frequency, max_frequency) to show in the plot.
        Note that these are only approximate since the data may
        not contain those exact frequencies. However, the
        plot is guaranteed not to show anything < min_frequency or
        >= max_frequency.
        Default is None, where all frequencies are shown.
    time_limits : list or None, optional
        If not None, a list consisting of the half-interval
        [min_time, max_time) to show in the plot. Note that these
        are only approximate since the data may not contain those
        exact time values. However, the plot is guaranteed not to
        show anything < min_time or >= max_time.
        Default is None, where all time points are shown.
    zscore : boolean, optional
        Whether to zscore the data before visualizing. zscoring
        is applied before restricting the display according to
        'freq_limits' and 'time_limits'
    zscore axis : None or int, optional
        The axis of s over which to apply zscoring. If None, zscoring
        is applied over the entire array s.
        Default is None.
    n_workers : integer, optional
        Number of parallel computations. Only used if 'zscore' is True.
        Default is the total number of CPUs (which may be virtual).
    timescale : string, optional
        The time scale to use on the plot's x-axis. Can be
        'milliseconds', seconds', 'minutes', or 'hours'.
        Default is 'seconds'
    relative_time : boolean, optional
        Whether the time axis shown on the plot will be relative
        Default is False
    center_time : boolean, optional
        Whether the time axis is centered around 0. This option
        is only available if 'relative_time' is set to True
        Default is False
    colorbar : boolean, optional
        Whether to show colorbar or not.
        Default is True.
    plot_type : {'pcolormesh', 'contourf', 'quadmesh'}, optional
        Type of plot to show. Note that the 'quadmesh' option uses
        hvplot as the backend.
        Default is 'pcolormesh'
    fig : matplotlib figure
        Used if 'colorbar' is True and 'plot_type' is not 'quadmesh'
    ax : matplotlib axes
        If None, new axes will be generated. Note that this argument
        is ignored if plot_type=='quadmesh'
    **kwargs: optional
        Other keyword arguments. Passed directly to pcolormesh(),
        contourf(), or quadmesh().
        
    Returns
    -------
    If plot_type is 'pcolormesh' or 'contourf':
        fig, ax : tuple consisting of (matplotlib figure, matplotlib axes)
    Otherwise:
        plot : bokeh plot handle
    
    """
    
    if freq_limits is not None:
        freq_slices = _get_frequency_slices(f, freq_limits[0], freq_limits[1])
    else:
        freq_slices = slice(None, None, None)
    
    if time_limits is not None:
        time_slices = _get_time_slices(t, time_limits[0], time_limites[1])
    else:
        time_slices = slice(None, None, None)
    
    tvec = t[time_slices]
    if timescale == 'milliseconds':
        tvec = tvec * 1000
        xlabel = "Time (msec)"
    elif timescale == 'seconds':
        xlabel = "Time (sec)"
    elif timescale == 'minutes':
        tvec = tvec / 60
        xlabel = "Time (min)"
    else:
        tvec = tvec / 3600
        xlabel = "Time (hr)"

    if relative_time:
        tvec = tvec - tvec[0]
        if center_time:
            a = (tvec[0] + tvec[-1]) / 2
            tvec = tvec - a
    
    fvec = f[freq_slices]
    df = np.diff(fvec)[0]
    dt = np.diff(tvec)[0]
    fbin_edges = np.linspace(fvec[0] - df/2, fvec[-1] + df/2, num=len(fvec)+1)
    tbin_edges = np.linspace(tvec[0] - dt/2, tvec[-1] + dt/2, num=len(tvec)+1)
    
    if zscore:
        # data may be large. use dask for computation
        dask_data = da.from_array(s)
        if zscore_axis is None:
            dask_data = (dask_data - dask_data.mean()) / dask_data.std()
        else:
            dask_data = (
                (dask_data - dask_data.mean(axis=zscore_axis, keepdims=True)) /
                 dask_data.std(axis=zscore_axis, keepdims=True)
            )
        data = dask_data[freq_slices, time_slices].compute(num_workers=n_workers)
    else:
        data = s[freq_slices, time_slices]
            
    if ax is None and plot_type != 'quadmesh':
        fig, ax = plt.subplots(1, 1)
        
    if colorbar:
        if (fig is None and ax is not None) or (fig is not None and ax is None): 
            raise ValueError(
                "Both 'fig' and 'ax' must be passed in if either is specified")
        
    if plot_type == 'pcolormesh':
        _set_matplotlib(True)
        im = ax.pcolormesh(tbin_edges, fbin_edges, data, **kwargs)
        if colorbar:
            fig.colorbar(im, ax=ax)
        ax.set_title("Spectrogram")
        ax.set_ylabel("Frequency")
        ax.set_xlabel(xlabel)
        return fig, ax
    elif plot_type == 'contourf':
        _set_matplotlib(True)
        im = ax.contourf(tvec, fvec, data, **kwargs)
        if colorbar:
            fig.colorbar(im, ax=ax)
        ax.set_title("Spectrogram")
        ax.set_ylabel("Frequency")
        ax.set_xlabel(xlabel)
        return fig, ax
    elif plot_type == 'quadmesh':
        _set_matplotlib(False)
        xa = xr.DataArray(
            data,
            dims=['Frequency', xlabel],
            coords={'Frequency':fvec, xlabel:tvec})
        plot = xa.hvplot(
            x=xlabel, y='Frequency', title="Spectrogram", colorbar=colorbar, **kwargs)
        return plot


def plot_wavelet_spectrogram(s, f, t, *, kind='amplitude', freq_limits=None, time_limits=None,
                             zscore=False, zscore_axis=None, n_workers=cpu_count(),
                             timescale='seconds', relative_time=False, center_time=False,
                             colorbar=True, plot_type='pcolormesh', fig=None,
                             ax=None, **kwargs):
    """
    Plot a wavelet based spectrogram. Note that the spectrogram
    data need not fit into memory but the data used for the
    plot should.

    Parameters
    ----------
    s : np.ndarray, with shape (n_freqs, n_timepoints)
        The spectrogram. Should be real-valued.
    f : np.ndarray, with shape (n_freqs, )
        Spectrogram frequencies in Hz. Must be uniformly
        spaced and strictly increasing or strictly
        decreasing.
    t : np.ndarray, with shape (n_timepoints, )
        Midpoints of each spectrogram time window in seconds.
        Must be uniformly spaced and strictly increasing
        or strictly decreasing.
    kind : {'amplitude', 'power'}, optional
        Display the data using the wavelet coefficient amplitudes
        or power, depending on what 'kind' is.
        Default is 'amplitude'.
    freq_limits : list or None, optional
        If not None, a list consisting of the half-open interval
        [min_frequency, max_frequency) to show in the plot.
        Note that these are only approximate since the data may
        not contain those exact frequencies. However, the
        plot is guaranteed not to show anything < min_frequency or
        >= max_frequency.
        Default is None, where all frequencies are shown.
    time_limits : list or None, optional
        If not None, a list consisting of the half-interval
        [min_time, max_time) to show in the plot. Note that these
        are only approximate since the data may not contain those
        exact time values. However, the plot is guaranteed not to
        show anything < min_time or >= max_time.
        Default is None, where all time points are shown.
    zscore : boolean, optional
        Whether to zscore the data before visualizing. zscoring
        is applied before restricting the display according to
        'freq_limits' and 'time_limits'
    zscore axis : None or int, optional
        The axis of s over which to apply zscoring. If None, zscoring
        is applied over the entire array s.
        Default is None.
    n_workers : integer, optional
        Number of parallel computations. Only used if 'zscore' is True.
        Default is the total number of CPUs (which may be virtual).
    timescale : string, optional
        The time scale to use on the plot's x-axis. Can be
        'milliseconds', seconds', 'minutes', or 'hours'.
        Default is 'seconds'
    relative_time : boolean, optional
        Whether the time axis shown on the plot will be relative
        Default is False
    center_time : boolean, optional
        Whether the time axis is centered around 0. This option
        is only available if 'relative_time' is set to True
        Default is False
    colorbar : boolean, optional
        Whether to show colorbar or not.
        Default is True.
    plot_type : {'pcolormesh', 'contourf', 'quadmesh'}, optional
        Type of plot to show. Note that the 'quadmesh' option uses
        hvplot as the backend.
        Default is 'pcolormesh'
    fig : matplotlib figure
        Used if 'colorbar' is True and 'plot_type' is not 'quadmesh'
    ax : matplotlib axes
        If None, new axes will be generated. Note that this argument
        is ignored if plot_type=='quadmesh'
    **kwargs: optional
        Other keyword arguments. Passed directly to pcolormesh(),
        contourf(), or quadmesh().
        
    Returns
    -------
    If plot_type is 'pcolormesh' or 'contourf':
        fig, ax : tuple consisting of (matplotlib figure, matplotlib axes)
    Otherwise:
        plot : bokeh plot handle
    
    """
    if not np.iscomplexobj(s):
        raise TypeError("Expected input data to be complex")
    
    if kind == 'amplitude':
        title_str = "Amplitude"
    elif kind == 'power':
        title_str = "Power"
    
    if freq_limits is not None:
        freq_slices = _get_frequency_slices(f, freq_limits[0], freq_limits[1])
    else:
        freq_slices = slice(None, None, None)
    
    if time_limits is not None:
        time_slices = _get_time_slices(t, time_limits[0], time_limites[1])
    else:
        time_slices = slice(None, None, None)

    tvec = t[time_slices]
    if timescale == 'milliseconds':
        tvec = tvec * 1000
        xlabel = "Time (msec)"
    elif timescale == 'seconds':
        xlabel = "Time (sec)"
    elif timescale == 'minutes':
        tvec = tvec / 60
        xlabel = "Time (min)"
    else:
        tvec = tvec / 3600
        xlabel = "Time (hr)"

    if relative_time:
        tvec = tvec - tvec[0]
        if center_time:
            a = (tvec[0] + tvec[-1]) / 2
            tvec = tvec - a
    
    fvec = f[freq_slices]
    
    if zscore:
        # data may be large. use dask for computation
        dask_data = da.absolute(da.from_array(s))
        if kind == 'power':
            dask_data = da.square(dask_data)
        
        if zscore_axis is None:
            dask_data = (dask_data - dask_data.mean()) / dask_data.std()
        else:
            dask_data = (
                (dask_data - dask_data.mean(axis=zscore_axis, keepdims=True)) /
                 dask_data.std(axis=zscore_axis, keepdims=True)
            )
        data = dask_data[freq_slices, time_slices].compute(num_workers=n_workers)
    else:
        data = np.abs(s[freq_slices, time_slices])
        if kind == 'power':
            data = data**2
            
    if ax is None and plot_type != 'quadmesh':
        fig, ax = plt.subplots(1, 1)
        
    if colorbar:
        if (fig is None and ax is not None) or (fig is not None and ax is None): 
            raise ValueError(
                "Both 'fig' and 'ax' must be passed in if either is specified")
        
    if plot_type == 'pcolormesh':
        _set_matplotlib(True)
        im = ax.pcolormesh(tvec, fvec, data, **kwargs)
        if colorbar:
            fig.colorbar(im, ax=ax)
        ax.set_title(title_str)
        ax.set_xlabel(xlabel)
        ax.set_ylabel("Frequency")
        return fig, ax
    elif plot_type == 'contourf':
        _set_matplotlib(True)
        im = ax.contourf(tvec, fvec, data, **kwargs)
        if colorbar:
            fig.colorbar(im, ax=ax)
        ax.set_title(title_str)
        ax.set_xlabel(xlabel)
        ax.set_ylabel("Frequency")
        return fig, ax
    elif plot_type == 'quadmesh':
        _set_matplotlib(False)
        xa = xr.DataArray(
            data,
            dims=['Frequency', xlabel],
            coords={'Frequency':fvec, xlabel:tvec})
        plot = xa.hvplot.quadmesh(
            x=xlabel, y='Frequency', title=title_str,
            colorbar=colorbar, **kwargs)
        return plot
    
def plot_frequency_response(b, a=1, *, fs=1, use_matplotlib=True, ax=None, **kwargs):
    """
    Plots frequency response (magnitude and phase) of a transfer function
    
    Parameters
    ----------
    b : float or np.ndarray with one dimension
        Numerator
    a : float or np.ndarray with one dimension, optional
        Denominator. 
        Default is 1.
    fs : float, optional
        Sampling rate in Hz.
    use_matplotlib : boolean, optional
        Whether to use matplotlib plotting backend.
        If True, produces a static plot. If False, produces
        an interactive plot with hvplot.
        Default is True.
    ax : matplotlib axes, optional
        Only used if 'use_matplotlib' is True.
        Default is None (axes are generated)
    **kwargs : optional
        Keyword arguments passed directly to matplotlib
        plot() or hvplot.line(), depending on the value
        of 'use_matplotlib'
        
    Returns
    -------
    If 'use_matplotlib' is True:
        ax : matplotlib axes
    Otherwise:
        layout : hvplot layout
        
    """

    w, h = sig.freqz(b, a, fs=fs)
    mag = 20*np.log10(np.abs(h))
    phase = np.angle(h)
    
    if use_matplotlib:
        _set_matplotlib(True)
        
        if ax is None:
            fig, ax = plt.subplots(2, 1)

        ax[0].plot(w, mag)
        ax[0].set_title("Magnitude Response")
        ax[0].set_xlabel("Frequency (Hz)")
        ax[0].set_ylabel("Magnitude (dB)")
        
        ax[1].plot(w, phase)
        ax[1].set_title("Phase Response")
        ax[1].set_xlabel("Frequency (Hz)")
        ax[1].set_ylabel("Angle (radians)")
        
        plt.tight_layout()
        
        return ax
    else:
        _set_matplotlib(False)
        
        magnitude_response = xr.DataArray(
            mag, dims=['Frequency'], coords={'Frequency':w}, name='Magnitude (dB)')
        phase_response = xr.DataArray(
            phase, dims=['Frequency'], coords={'Frequency':w}, name='Phase (radians)')
        
        return (
            magnitude_response.hvplot.line(title='Magnitude Response', x='Frequency') + 
            phase_response.hvplot.line(title='Phase Response', x='Frequency')
            ).cols(1)

def _get_frequency_slices(freqs, flow, fhigh):
    if freqs[0] < freqs[-1]: # ascending order
        ind1 = np.argwhere(freqs >= flow).squeeze().min()
        ind2 = np.argwhere(freqs <= fhigh).squeeze().max()
        return slice(ind1, ind2)
    else:
        ind1 = np.argwhere(freqs <= fhigh).squeeze().min()
        ind2 = np.argwhere(freqs >= flow).squeeze().max()
        return slice(ind1, ind2)
    
def _get_time_slices(times, tlow, thigh):
    if times[0] < times[-1]:
        ind1 = np.argwhere(times >= tlow).squeeze().min()
        ind2 = np.argwhere(times <= thigh).squeeze().max()
        return slice(ind1, ind2)
    else:
        ind1 = np.argwhere(times <= thigh).squeeze().min()
        ind2 = np.argwhere(times >= tlow).squeeze().max()
        return slice(ind1, ind2)
    
def _set_matplotlib(matplotlib):
    if matplotlib:
        hv.extension('matplotlib')
    else:
        hv.extension('bokeh')
```
