## Supplementary material for "GhostiPy: an efficient signal processing and spectral analysis toolbox for large data": Source code: cwt.py

```
import logging
import pyfftw
import dask
import warnings
import numpy as np

import time
import traceback

from multiprocessing import cpu_count
from dask.diagnostics import ProgressBar
from ghostipy.utils import (hz_to_normalized_rad, normalized_rad_to_hz)
from ghostipy.spectral.wavelets import (Wavelet, MorseWavelet, AmorWavelet, BumpWavelet)

__all__ = ['cwt']

def cwt(data, *, fs=1, timestamps=None, wavelet=MorseWavelet(gamma=3, beta=20),
                freq_limits=None, freqs=None, voices_per_octave=10,
                n_workers=cpu_count(), verbose=False, method='full',
                derivative=False, remove_mean=False, boundary='mirror',
                coi_threshold=1/(np.e**2), describe_dims=False,
                cwt_out=None):
    """Computes the continuous wavelet transform.

    Parameters
    ----------
    data : np.ndarray, with shape (n_timepoints, )
        Data with which to compute the CWT
    fs : float
        Sampling rate of the data in Hz.
    timestamps : np.ndarray, with shape (n_timepoints, ) optional
        Timestamps corresponding to the data, in seconds.
        If None, they will be computed automatically based on the
        assumption that all the data are one contiguous block, and
        the units will be in seconds.
    wavelet : ghostipy.wavelet
        Type of wavelet to use for the transform.
        Default is a Morse wavelet with beta=3 and gamma=20.
    freq_limits : list, optional
        List of [lower_bound, upper_bound] for frequencies to use,
        in units of Hz. Note that a reference set of frequencies
        is generated on the shortest segment of data since that
        determines the lowest frequency that can be used. If the
        bounds specified by 'freq_limits' are outside the bounds
        determined by the reference set, 'freq_limits' will be
        adjusted to be within the bounds of the reference set.
    freqs : array-like, optional
        Frequencies to analyze, in units of Hz.
        Note that a reference set of frequencies is computed on the
        shortest segment of data since that determines the lowest
        frequency that can be used. If any frequencies specified in
        'freqs' are outside the bounds determined by the reference
        set, 'freqs' will be adjusted such that all frequencies in
        'freqs' will be within those bounds.
    voices_per_octave : int, optional
        Number of wavelet frequencies per octave. Note that this
        parameter is not used if frequencies were already specified
        by the 'freqs' option.
        Default is 10.
    n_workers : integer, optional
        Number of parallel jobs to use.
        Default is the total number of CPUs (which may be virtual).
    verbose : boolean, optional
        Whether to print messages displaying this function's progress.
        Default is False.
    method: {'full', 'ola'}, optional
        Which algorithm to use for computing the CWT. 'ola' will give
        superior performance for long lengths of data.
    derivative: boolean, optional
        Whether to compute the derivative of the CWT.
        Default is False.
    remove_mean: boolean, optional
        Whether to remove the mean of the data before computing the CWT.
        Default is False.
    boundary: {'mirror', 'zeros', 'periodic'}, optional
        To handle boundaries, the data are extended before computing the CWT
        according to the following options:
        'mirror' : Mirror/reflect the data at each end
        'zeros': Add zeros at each end
        'periodic': Treat the data as periodic
        Note that regardless of the boundary method, the CWT should not be
        treated as reliable within the cone of influence.
        Default is 'mirror'.
    coi_threshold : float, optional
        The value C that determines the wavelet's cone of influence. The
        maximum value P of the wavelet's power autocorrelation is taken
        in the time domain. Then the cone of influence is given by the
        region where the power autocorrelation is above C*P. Default value
        for C is e^(-2).
    describe_dims : boolean, optional
        Whether to return the expected shape and dtype of the output and
        return immediately (no CWT is computed). This option is
        useful for out-of-core computation. While the expected shape
        should not be changed, the dtype is only suggested, e.g. it is
        acceptable to use a lower precision dtype (such as complex64 instead
        of complex128 to save space)
        Default is False.
    cwt_out: array-like, optional
        If specified, the CWT output coefficients will be stored here.
        Useful if the output is too large to fit into memory and must instead
        be saved to an array stored on disk.
    

    Returns
    -------
    If 'describe_dims' is True:
        shape, dtype : tuple
            Expected output array shape and dtype
    Otherwise:
        coefs : np.ndarray, with shape (n_scales, n_timepoints)
            Calculated continuous wavelet coefficients. Note that the scale
            dimension is ordered by increasing wavelet scale, which corresponds
            to decreasing frequency.
        scales : np.ndarray, with shape (n_scales, )
            Wavelet scales for which CWT was calculated.
        frequencies : np.ndarray, with shape (n_frequencies, )
            If a sampling rate is given, these will be the frequency
            corresponding to each wavelet scale.
        timestamps : np.array, with shape (n_timepoints, )
            If timestamps were not specified, these are the timestamps
            calculated for data samples. Pass through of input 
            timestamps otherwise.
        cois : np.ndarray, with shape (n_cois, )
            Cones of influence for each wavelet scale.
    """

    t0 = time.time()
    if verbose:
        print("Using maximum of {} workers".format(n_workers))

    if not isinstance(wavelet, Wavelet):
        raise TypeError("Supplied wavelet must inherit from a ghostipy.Wavelet type")

    if freqs is not None and freq_limits is not None:
        raise ValueError("'freqs' and 'freq_limits' cannot both be used at the"
                            " same time. Either specify one or the other, or"
                            " leave both as unspecified")

    if freqs is not None and voices_per_octave is not None:
        raise ValueError("'freqs' and 'voices_per_octave' cannot both be used"
                            " at the same time. Either specify one or the other,"
                            " or leave both as unspecified")

    if method not in ('full', 'ola'):
        raise ValueError(f"Invalid method {method}")

    N = data.shape[0]

    if verbose:
        print("Determining smallest scale...")
    ref_scale, ref_coi = wavelet.reference_coi(threshold=coi_threshold)
    max_scale = N / ref_coi * ref_scale

    w_ref_low = wavelet.scale_to_freq(max_scale).squeeze()
    w_ref_high = np.pi
    if verbose:
        print(
            f"Smallest reference frequency: {normalized_rad_to_hz(w_ref_low, fs):0.4f} Hz")

    if freqs is not None:
        # just in case user didn't pass in sorted
        # frequencies after all
        freqs = np.sort(freqs)
        ws = hz_to_normalized_rad(freqs, fs)
        w_low = ws[0]
        w_high = ws[-1]
        if w_low < w_ref_low:
            warnings.warn(
                f"Warning: Lower frequency limit of {freq_limits[0]} is less than the smallest "
                f"recommended frequency of {normalized_rad_to_hz(w_ref_low, fs):0.4f} Hz")
        if w_high > w_ref_high:
            warnings.warn(
                f"Warning: Upper frequency limit of {freq_limits[1]} is greater than the largest "
                f"recommended frequency of {normalized_rad_to_hz(w_ref_high, fs):0.4f} Hz")
    elif freq_limits is not None:
        # just in case user didn't pass in limits as [lower_bound, upper_bound]
        freq_limits = np.sort(freq_limits)
        w_low = hz_to_normalized_rad(freq_limits[0], fs)
        w_high = hz_to_normalized_rad(freq_limits[1], fs)
        if w_low < w_ref_low:
            warnings.warn(
                f"Lower frequency limit of {freq_limits[0]} is less than the smallest "
                f"recommended frequency of {normalized_rad_to_hz(w_ref_low, fs):0.4f} Hz")
        if w_high > w_ref_high:
            warnings.warn(
                f"Upper frequency limit of {freq_limits[1]} is greater than the largest "
                f"recommended frequency of {normalized_rad_to_hz(w_ref_high, fs):0.4f} Hz")
    else:
        w_low = w_ref_low
        w_high = w_ref_high

    if freqs is None:
        n_octaves = np.log2(w_high / w_low)
        j = np.arange(n_octaves * voices_per_octave)
        ws = w_high / 2**(j/voices_per_octave)

    scales = wavelet.freq_to_scale(ws)
    cois = wavelet.coi(scales, ref_scale, ref_coi)


    if remove_mean:
        # Don't do in place here, even though it saves memory,
        # as that would mutate the original data
        data = data - data.mean()
    
    extend_len = int(np.ceil(np.max(cois)))
    if extend_len > N:
        warnings.warn(f"Cannot add {extend_len} points to satisfy requested"
                f" boundary policy. Shorting this value to data length {N}")
        extend_len = N
    
    if boundary == 'mirror':
        data = np.hstack((np.flip(data[:extend_len]), data, np.flip(data[-extend_len:])))
    elif boundary == 'zeros':
        data = np.hstack((np.zeros(extend_len), data, np.zeros(extend_len)))
    elif boundary == 'periodic':
        data = np.hstack((data[-extend_len:], data, data[:extend_len]))
    else:
        extend_len = 0

    # Set up array as C contiguous since we will be iterating row-by-row
    n_bits = len(scales) * data.shape[0] * 16
    if verbose:
        print(f"Output space requirement: {n_bits/1e9} GB = {n_bits/(1024**3)} GiB")

    output_shape = (scales.shape[0], N)
    dtype = '<c16'
    if describe_dims:
        if verbose:
            print("Calculating output array sizes. Skipping transform computation")
            print(f"Output array with 'derivative' {derivative}"
                  f" should have shape {output_shape} with dtype {dtype}")
        return output_shape, dtype

    if cwt_out is not None:
        if verbose:
            print("Using passed-in output array")
        if cwt_out.shape != output_shape:
            raise ValueError(
                f"Provided output array has shape {coefs.shape}"
                f" but needs shape {output_shape}")
        coefs = cwt_out
    else:
        if verbose:
            print("Allocating output array")
        coefs = pyfftw.zeros_aligned(output_shape, dtype='complex128')

    ######################################################################
    # Set up CWT parallel computation

    task_list = []
    if method == 'ola':
        for ii in range(scales.shape[0]):
            task = dask.delayed(_cwt_ola_fftw)(
                data,
                wavelet,
                scales[ii],
                derivative,
                coefs,
                ii,
                extend_len,
                1,
                cois[-1])

            task_list.append(task)
        
    elif method == 'full':
        if verbose:
            print("Computing FFT of data")

        data_fft = pyfftw.zeros_aligned(data.shape[0], dtype='complex128')
        fft_sig = pyfftw.FFTW(data_fft, data_fft,
                        direction='FFTW_FORWARD',
                        flags=['FFTW_ESTIMATE'],
                        threads=n_workers)
        data_fft[:] = data
        fft_sig()
        omega = np.fft.fftfreq(data.shape[0]) * 2 * np.pi
        for ii in range(scales.shape[0]):
            task = dask.delayed(_cwt_full_fftw)(
                data_fft,
                wavelet,
                scales[ii],
                derivative,
                coefs,
                ii,
                extend_len,
                1,
                omega)

            task_list.append(task)

    if verbose:
        print("Computing CWT coefficients")
        with ProgressBar():
            dask.compute(task_list, num_workers=n_workers)
    else:
        dask.compute(task_list, num_workers=n_workers)


    if verbose:
        print('CWT total elapsed time: {} seconds'.format(time.time() - t0))

    if timestamps is None:
        timestamps = np.arange(N) / fs
    
    return coefs, scales, normalized_rad_to_hz(ws, fs), timestamps, cois

def _cwt_ola_fftw(data, wavelet, scale, derivative, cwt, ii,
                  extend_len, threads, coi):

    """
    Parameters
    ----------
    data : numpy.ndarray
        Input data

    wavelet : of type ghost.Wavelet
        Wavelet to use for the CWT

    Returns
    -------
    cwt : numpy.ndarray
        The (complex) continuous wavelet transform

    Notes
    -----
    This function is a specialized method intended specifically for
    the CWT and was written with the following principles/motivations
    in mind:

    (1) We want to compute the wavelet transform as numerically exact as
    possible. For most cases, we can compute a wavelet's time-domain
    representation over its footprint/effective support and use overlap-add
    for efficient chunked convolution. However, this strategy can be
    problematic for analytic wavelets. If the wavelet's frequency response
    is not zero at the input data's Nyquist frequency, there will be
    Gibbs' phenomenon when taking the DFT of the wavelet's time-domain
    representation. Then the wavelet is no longer numerically analytic.

    (2) We therefore compute the wavelet directly in the frequency domain,
    with a length longer than its footprint/effective support. Then we take
    overlapping data chunks, similar to the overlap-save method. However,
    unlike overlap-save our method discards BOTH edges of each chunk to
    deal with boundary effects. Excluding the edges at the very ends of the
    computed wavelet transform, this result is identical to that obtained
    if the transform were computed over the data in one long chunk. Thus, 
    this method is an efficient way to calculate the wavelet transform on
    chunks of data while handling potential numerical issues associated with
    analytic wavelets.
    """
    M = int(np.ceil(coi))
    N = 65536 # good default FFT length for most frequencies of interest
    while N < 10 * M: # make sure sufficiently long
        N *= 4

    start, stop = 0, len(data)
    buf_starts = np.arange(start, stop, N-2*M)

    psif = np.zeros((1, N), dtype='<c16')
    omega = np.fft.fftfreq(N) * 2 * np.pi

    wavelet.freq_domain_numba(
        omega,
        np.atleast_1d(scale),
        psif,
        derivative=derivative)

    if not wavelet.is_analytic:
        psif = psif.conj()

    ########################################################################
    x = pyfftw.zeros_aligned(N, dtype='complex128')
    # using powers of 4 so FFTW_ESTIMATE should suffice
    # in place transform to save memory
    fft_sig = pyfftw.FFTW( 
        x, x,
        direction='FFTW_FORWARD',
        flags=['FFTW_ESTIMATE'],
        threads=threads)

    y = pyfftw.zeros_aligned(N, dtype='complex128')
    y[:] = psif

    conv_res = pyfftw.zeros_aligned(N, dtype='complex128')
    fft_conv_inv = pyfftw.FFTW(
        conv_res, conv_res,
        direction='FFTW_BACKWARD', 
        flags=['FFTW_ESTIMATE'], 
        threads=threads)

    #######################################################################
    first_block_to_check, first_offset = divmod(extend_len, N - 2*M)
    last_block_to_check, last_offset = divmod(stop - extend_len, N - 2*M)
#     print(f'Need to check blocks {first_block_to_check} and {last_block_to_check},'
#           f' {buf_starts.shape[0]} total blocks')
    outarray_marker = 0
    for block_ind, buf_start in enumerate(buf_starts):

        if buf_start - M < 0:
            first_segment = data[0:buf_start]
            length = len(first_segment)
            x[:M-length] = 0
            x[M-length:M] = first_segment
        else:
            first_segment = data[buf_start - M:buf_start]
            if len(first_segment) < M:
                x[:len(first_segment)] = first_segment
                x[len(first_segment):M] = 0
            else:
                x[:M] = first_segment

        second_segment = data[buf_start:buf_start + N - M]
        x[M:M+len(second_segment)] = second_segment
        x[M+len(second_segment):] = 0

        fft_sig(normalise_idft=True)
        conv_res[:] = x * y
        fft_conv_inv(normalise_idft=True)

        cwt_chunk = conv_res[M:N-M]

        if block_ind == first_block_to_check and block_ind == last_block_to_check:
            n_samples = last_offset - first_offset
            cwt[ii, :n_samples] = cwt_chunk[first_offset:last_offset]
            outarray_marker += n_samples
        elif block_ind == first_block_to_check:
            n_samples = len(cwt_chunk) - first_offset
            cwt[ii, :n_samples] = cwt_chunk[first_offset:]
            outarray_marker += n_samples
        elif block_ind == last_block_to_check:
            n_samples = last_offset
            cwt[ii, outarray_marker:outarray_marker+n_samples] = cwt_chunk[:n_samples]
            outarray_marker += n_samples
        elif block_ind > first_block_to_check and block_ind < last_block_to_check:
            n_samples = len(cwt_chunk)
            cwt[ii, outarray_marker:outarray_marker+n_samples] = cwt_chunk
            outarray_marker += n_samples
        else:
            pass

    return

def _cwt_full_fftw(data_fft, wavelet, scale, derivative, cwt, ii,
                   extend_len, threads, omega):

    out = pyfftw.zeros_aligned((1, len(data_fft)), dtype='complex128')

    fft_conv_inv = pyfftw.FFTW(
        out, out,
        axes=(-1, ),
        direction='FFTW_BACKWARD',
        flags=['FFTW_ESTIMATE'],
        threads=threads)

    wavelet.freq_domain_numba(
        omega,
        np.atleast_1d(scale),
        out,
        derivative=derivative)

    if not wavelet.is_analytic:
        out[:] = out.conj()

    out *= data_fft
    fft_conv_inv(normalise_idft=True)
    cwt[ii:ii+1] = out[:, extend_len:len(data_fft)-extend_len]

    return
```
