## Supplementary material for "GhostiPy: an efficient signal processing and spectral analysis toolbox for large data": Source code: mtm.py

```
import pyfftw
import numpy as np
from numpy.lib.stride_tricks import as_strided
from multiprocessing import cpu_count
from scipy.signal.windows import dpss

__all__ = ['get_tapers',
           'mtm_spectrum',
           'mtm_spectrogram']

def get_tapers(N, bandwidth, *, fs=1, min_lambda=0.95, n_tapers=None):
    """
    Compute tapers and associated energy concentrations for the Thomson
    multitaper method

    Parameters
    ----------
    N : int
        Length of taper
    bandwidth : float
        Bandwidth of taper, in Hz
    fs : float, optional
        Sampling rate, in Hz.
        Default is 1 Hz.
    min_lambda : float, optional
        Minimum energy concentration that each taper must satisfy.
        Default is 0.95.
    n_tapers : int, optional
        Number of tapers to compute
        Default is to use all tapers that satisfied 'min_lambda'.
    
    Returns
    -------
    tapers : np.ndarray, with shape (n_tapers, N)
    lambdas : np.ndarray, with shape (n_tapers, )
        Energy concentrations for each taper

    """
    
    NW = bandwidth * N / fs 
    K = int(np.ceil(2*NW)) - 1
    if K < 1:
        raise ValueError(
            f"Not enough tapers, with 'NW' of {NW}. Increase the bandwidth or "
            "use more data points")
        
    tapers, lambdas = dpss(N, NW, Kmax=K, norm=2, return_ratios=True)
    mask = lambdas > min_lambda
    if not np.sum(mask) > 0:
        raise ValueError(
            "None of the tapers satisfied the minimum energy concentration"
            f" criteria of {min_lambda}")
    tapers = tapers[mask]
    lambdas = lambdas[mask]

    if n_tapers is not None:
        if n_tapers > tapers.shape[0]:
            raise ValueError(
                f"'n_tapers' of {n_tapers} is greater than the {tapers.shape[0]}"
                f" that satisfied the minimum energy concentration criteria of {min_lambda}")
        tapers = tapers[:n_tapers]
        lambdas = lambdas[:n_tapers]
    
    return tapers, lambdas

def mtm_spectrum(data, bandwidth, *, fs=1, min_lambda=0.95, n_tapers=None,
                 remove_mean=False, nfft=None, n_fft_threads=cpu_count()):

    """
    Computes the spectrum using Thomson's multitaper method.

    Parameters
    ----------
    data : np.ndarray, with shape (T, )
        Input data
    bandwidth : float
        Bandwidth of taper, in Hz
    fs : float, optional
        Sampling rate, in Hz.
        Default is 1 Hz.
    min_lambda : float, optional
        Minimum energy concentration that each taper must satisfy.
        Default is 0.95.
    n_tapers : int, optional
        Number of tapers to compute
        Default is to use all tapers that satisfied 'min_lambda'.
    remove_mean : boolean, optional
        Whether to remove the mean of the data before computing the
        MTM spectrum.
        Default is False.
    nfft : int, optional
        How many FFT points to use for the spectrum.
        Default is the same as the length of the input data.
    n_fft_threads : int, optional
        Number of threads to use for the FFT.
        Default is the number of CPUs (which may be virtual).
    
    Returns
    -------
    mt_sdf : np.ndarray, with shape (nfft, )
        The multitapered power spectral density
    freqs : np.ndarray, with shape (nfft, )
        The corresponding frequencies for the mt PSD, in Hz.

    """

    N = data.shape[0]
    tapers, lambdas = get_tapers(
        N, bandwidth, fs=fs,
        n_tapers=n_tapers,
        min_lambda=min_lambda)
    n_tapers = tapers.shape[0]
    
    if nfft is None:
        nfft = N
        
    if remove_mean:
        data = data - data.mean()

    if np.isrealobj(data):
        M = nfft // 2 + 1

        xtd = pyfftw.zeros_aligned(
            (n_tapers, nfft),
            dtype='float64')
        xfd = pyfftw.zeros_aligned(
            (n_tapers, M),
            dtype='complex128')
        fft_sig = pyfftw.FFTW( 
            xtd, xfd,
            axes=(1, ),
            direction='FFTW_FORWARD',
            flags=['FFTW_ESTIMATE'],
            threads=n_fft_threads,
            planning_timelimit=0)
        
        xtd[:, :N] = tapers * data
        xtd[:, N:] = 0
        fft_sig(normalise_idft=True)
        #assert np.allclose(xfd, np.fft.rfft(tapers * data, n=nfft))
        #xfd = np.fft.rfft(tapers * data, n=nfft)
        
        sdfs = (xfd.real**2 + xfd.imag**2) / fs
        
        if nfft % 2 == 0:
            sdfs[:, 1:-1] *= 2
        else:
            sdfs[:, 1:] *= 2

        freqs = np.fft.rfftfreq(nfft, d=1/fs)
    else:
        # can use an in-place transform here
        x = pyfftw.zeros_aligned((n_tapers, nfft), dtype='complex128')
        fft_sig = pyfftw.FFTW( 
            x, x,
            axes=(1, ),
            direction='FFTW_FORWARD',
            flags=['FFTW_ESTIMATE'],
            threads=n_fft_threads,
            planning_timelimit=0)
        
        x[:, :N] = tapers * data
        x[:, N:] = 0
        fft_sig(normalise_idft=True)
        #assert np.allclose(xfd, np.fft.fft(tapers * data, n=nfft))

        sdfs = (x.real**2 + x.imag**2) / fs        
        freqs = np.fft.fftfreq(nfft, d=1/fs)

    mt_sdf = np.mean(sdfs, axis=0)
        
    return mt_sdf, freqs


def mtm_spectrogram(data, bandwidth, *, fs=1, timestamps=None, nperseg=None, noverlap=None,
                    n_tapers=None, min_lambda=0.95, remove_mean=False, nfft=None,
                    n_fft_threads=cpu_count()):

    """
    Computes the spectrogram using Thomson's multitaper method.

    Parameters
    ----------
    data : np.ndarray, with shape (T, )
        Input data
    bandwidth : float
        Bandwidth of taper, in Hz
    fs : float, optional
        Sampling rate, in Hz.
        Default is 1 Hz.
    timestamps : np.ndarray, with shape (T, ), optional
        Timestamps for the data. If not provided, they will be
        inferred using np.arange(len(data)) / fs
    nperseg : int, optional
        Number of samples to use for each segment/window.
        Default is 256.
    noverlap : int, optional
        Number of points to overlap between segments.
        Default is nperseg // 8.
    min_lambda : float, optional
        Minimum energy concentration that each taper must satisfy.
        Default is 0.95.
    n_tapers : int, optional
        Number of tapers to compute
        Default is to use all tapers that satisfied 'min_lambda'.
    remove_mean : boolean, optional
        Whether to remove the mean of the data before computing the
        MTM spectrum.
        Default is False.
    nfft : int, optional
        How many FFT points to use for each segment.
        Default is the value of 'nperseg'
    n_fft_threads : int, optional
        Number of threads to use for the FFT.
        Default is the number of CPUs (which may be virtual).
    
    Returns
    -------
    S : np.ndarray, with shape (n_freqs, n_timepoints)
        Multitapered spectrogram (units are power spectral density)
    f : np.narray, with shape (n_freqs, )
        Spectrogram frequencies
    t : np.ndarray, with shape (n_timepoints, )
        The midpoints of each segment/window.

    """
    
    N = data.shape[0]

    if timestamps is None:
        timestamps = np.arange(N) / fs

    if timestamps.shape[0] != N:
        raise ValueError(
            f"Expected timestamps to contain {N} elements but got {timestamps.shape[0]}")
        
    estimated_fs = 1.0/np.median(np.diff(timestamps))
    if np.abs((estimated_fs - fs)/fs) > 0.01:
        print("Warning: estimated fs and provided fs differ by more than 1%")

    if nperseg is None:
        nperseg = 256
    
    if noverlap is None:
        noverlap = nperseg // 8

    if noverlap >= nperseg:
        raise ValueError("noverlap must be less than {}".format(nperseg))
    
    if nfft is None:
        nfft = nperseg
    if nfft < nperseg:
        raise ValueError(f"'nfft' must be at least {nperseg}")
        
    if nperseg > N:
        raise ValueError(f"'nperseg' cannot be larger than the data size {N}")
        
    if not N > noverlap:
        raise ValueError(f"'noverlap' cannot be larger than {N-1}")
        
    if remove_mean:
        data = data - data.mean()

    tapers, lambdas = get_tapers(
        nperseg,
        bandwidth,
        fs=fs,
        n_tapers=n_tapers,
        min_lambda=min_lambda)
    n_tapers = tapers.shape[0]

    step = nperseg - noverlap
    shape = data.shape[:-1]+((data.shape[-1]-noverlap)//step, nperseg)
    strides = data.strides[:-1]+(step*data.strides[-1], data.strides[-1])
    data_strided = as_strided(
        data,
        shape=shape,
        strides=strides,
        writeable=False)
    
    n_segments = data_strided.shape[0]
    out_timestamps = np.mean(
        as_strided(
            timestamps,
            shape=shape,
            strides=strides,
            writeable=False),
            axis=1)
    
    if np.isrealobj(data):
        M = nfft // 2 + 1

        xtd = pyfftw.zeros_aligned(
            (n_tapers, n_segments, nfft),
            dtype='float64')
        xfd = pyfftw.zeros_aligned(
            (n_tapers, n_segments, M),
            dtype='complex128')
        fft_sig = pyfftw.FFTW( 
            xtd, xfd,
            axes=(2, ),
            direction='FFTW_FORWARD',
            flags=['FFTW_ESTIMATE'],
            threads=n_fft_threads,
            planning_timelimit=0)
        
        # (1, n_segments, nperseg) x (n_tapers, 1, nperseg)
        xtd[:, :, :N] = data_strided[None, :, :] * tapers[:, None, :]
        xtd[:, :, N:] = 0
        fft_sig(normalise_idft=True)
        #assert np.allclose(xfd, np.fft.rfft(data_strided[None, :, :] * tapers[:, None, :], n=nfft, axis=-1))
        #xfd = np.fft.rfft(tapers * data, n=nfft)
        
        spectrograms = (xfd.real**2 + xfd.imag**2) / fs
        
        if nfft % 2 == 0:
            spectrograms[:, :, 1:-1] *= 2
        else:
            spectrograms[:, :, 1:] *= 2

        freqs = np.fft.rfftfreq(nfft, d=1/fs)
    else:
        # can use an in-place transform here
        x = pyfftw.zeros_aligned(
            (n_tapers, n_segments, nfft),
            dtype='complex128')
        fft_sig = pyfftw.FFTW( 
            x, x,
            axes=(2, ),
            direction='FFTW_FORWARD',
            flags=['FFTW_ESTIMATE'],
            threads=n_fft_threads,
            planning_timelimit=0 )
        
        # (1, n_segments, nperseg) x (n_tapers, 1, nperseg)
        x[:, :, :N] = data_strided[None, :, :] * tapers[:, None, :]
        x[:, :, N:] = 0
        fft_sig(normalise_idft=True)
        #assert np.allclose(
        # xfd, np.fft.fft(data_strided[None, :, :] * tapers[:, None, :], n=nfft, axis=-1))

        spectrograms = (x.real**2 + x.imag**2) / fs      
        freqs = np.fft.fftfreq(nfft, d=1/fs)
    

    spectrogram = np.sum(lambdas[:, None, None] * spectrograms, axis=0) / np.sum(lambdas)
    assert np.all(np.isfinite(spectrogram))
    
    return spectrogram.T, freqs, out_timestamps
```
