## Supplementary material for "GhostiPy: an efficient signal processing and spectral analysis toolbox for large data": Source code: wavelets.py

```
import numpy as np
from abc import ABC, abstractmethod
from numba import njit
from scipy.signal import correlate

__all__ = ['Wavelet',
           'MorseWavelet',
           'AmorWavelet',
           'BumpWavelet']

def reference_coi(psifn, reference_scale, *, threshold=1/(np.e**2)):
    """
    Estimates a wavelet's cone of influence.

    Parameters
    ----------
    psifn : function
        Function to get the wavelet frequency domain representation
    reference_scale : float
        Scale at which 'psifn' should be evaluated
    threshold : float, optional
        The value C that determines the wavelet's cone of influence. The
        maximum value P of the wavelet's power autocorrelation is taken
        in the time domain. Then the cone of influence is given by the
        region where the power autocorrelation is above C*P. Default value
        for C is e^(-2).

    Returns
    -------
    reference_coi : float
        The COI for the passed-in 'reference_scale'

    """

    omega = np.fft.fftfreq(2**22) * 2 * np.pi
    psif = psifn(omega, reference_scale).squeeze()
    psit = np.fft.ifftshift(np.fft.ifft(psif))
    psit_power = psit.real**2 + psit.imag**2

    power_acs = correlate(psit_power, psit_power)
    maxind = np.argmax(power_acs)
    inds = np.argwhere(power_acs < threshold * np.max(power_acs)).squeeze()
    mask = inds > maxind
    inds = inds[mask]

    reference_coi = inds[0] - maxind

    return reference_coi

def coi(scales, reference_scale, reference_coi, *, fs=1):
    """
    Estimates a wavelet's cone of influence (in seconds)
    for requested scales, given a reference scale and
    reference COI

    Parameters
    ----------
    scales : np.ndarray, with shape (n_scales, )
        Array of scales for which to estimate the COI
    reference_scale : float
        The scale used as a reference
    reference_coi : float
        The COI used as a reference
    fs : float, optional
        Sampling rate.
        Default is 1, where the COI in seconds is identical
        to the COI in number of samples

    Returns
    -------
    cois : np.ndarray, with shape (n_scales, )
        The COIs for each value of 'scales'
    """
    
    scales = np.atleast_1d(scales)
    factors = scales / reference_scale

    return factors * reference_coi / fs

class Wavelet(ABC):
    """
    The abstract base class that all wavelets in this package inherit from.
    A custom wavelet should use this template if it is intended to be used
    for the cwt() and wsst() methods. Note that the built-in wavelets have
    been implemented as a bandpass filter bank with peak value 2 in the
    frequency domain.
    """

    def __init__(self):
        pass

    @abstractmethod
    def freq_domain(self, omega, scales, *, derivative=False):
        """
        Get the frequency domain representation of the wavelet

        Parameters
        ----------
        omega : np.ndarray, with shape (n_freqs, )
            Array of angular frequencies
        scales : np.narray, with shape (n_scales, )
            Array of scales to use
        derivative : boolean, optional
            If True, return the derivative of the wavelet

        Returns
        -------
        psif : np.ndarray, with shape (n_scales, n_freqs)
            The frequency domain representation given the
            passed-in 'scales'
        """
        pass

    @abstractmethod
    def freq_domain_numba(self, omega, scales, out, *, derivative=False):
        """
        Get the frequency domain representation of the wavelet using
        numba as the backend

        Parameters
        ----------
        omega : np.ndarray, with shape (n_freqs, )
            Array of angular frequencies
        scales : np.narray, with shape (n_scales, )
            Array of scales to use
        out : np.ndarray, with shape (n_scales, n_freqs)
            Output array to store the result
        derivative : boolean, optional
            If True, return the derivative of the wavelet

        Returns
        -------
        result : boolean
            True if successfully, False otherwise
        """
        pass

    @abstractmethod
    def freq_to_scale(self, freqs):
        """
        Map center frequency to scale

        Parameters
        ----------
        freqs : np.ndarray, with shape (n_freqs, )
            Array of frequencies. Units should be radians within
            the range [-pi, pi]

        Returns
        -------
        scales : np.ndarray, with shape (n_scales, )
            The scales corresponding to each frequency
        """
        pass

    @abstractmethod
    def scale_to_freq(self, scales):
        """
        Map scale to center frequency

        Parameters
        ----------
        scales : np.ndarray, with shape (n_scales, )
            Array of scales

        Returns
        -------
        freqs : np.ndarray, with shape (n_scales, )
            The center frequencies corresponding to each
            scale. Units are in radians.
        """
        pass

    @abstractmethod
    def reference_coi(self, *, threshold=1/(np.e**2)):
        """
        Get the COI for the base scale

        Parameters
        ----------
        threshold : float, optional
            The value C that determines the wavelet's cone of influence. The
            maximum value P of the wavelet's power autocorrelation is taken
            in the time domain. Then the cone of influence is given by the
            region where the power autocorrelation is above C*P. Default value
            for C is e^(-2).
        """
        pass

    @abstractmethod
    def coi(self, scales, reference_scale, reference_coi, *, fs=1):
        """
        Estimates a wavelet's cone of influence (in seconds)
        for requested scales, given a reference scale and
        reference COI

        Parameters
        ----------
        scales : np.ndarray, with shape (n_scales, )
            Array of scales for which to estimate the COI
        reference_scale : float
            The scale used as a reference
        reference_coi : float
            The COI used as a reference
        fs : float, optional
            Sampling rate.
            Default is 1, where the COI in seconds is identical
            to the COI in number of samples

        Returns
        -------
        cois : np.ndarray, with shape (n_scales, )
            The COIs for each value of 'scales'
        """
        pass

    @property
    @abstractmethod
    def admissibility_constant(self):
        """
        The admissibility constant (float)
        """
        pass

    @property
    @abstractmethod
    def is_analytic(self):
        """
        Whether or not a wavelet is analytic (boolean)
        """
        pass

@njit
def _morse_freq_domain(omega, scales, gamma, beta, out,
                       *, derivative=False):
    
    # out better be initialized to zeros!
    log_a = np.log(2) + (beta/gamma) * (1+np.log(gamma) - np.log(beta))

    H = np.zeros_like(omega)
    H[omega > 0] = 1

    for ii in range(scales.shape[0]):
        x = scales[ii] * omega
        log_psif = log_a + beta * np.log(np.abs(x)) - np.abs(x)**gamma
        out[ii] = np.exp(log_psif) * H
        if derivative:
            out[ii] *= 1j * omega

    return True

class MorseWavelet(Wavelet):
    
    def __init__(self, *, gamma=3, beta=20):
        
        super().__init__()
        self.gamma = gamma
        self.beta = beta
        self.wp = np.exp( (np.log(self.beta) - np.log(self.gamma)) / self.gamma )
        
    def freq_domain(self, omega, scales, *, derivative=False):
        
        gamma = self.gamma
        beta = self.beta

        scales = np.atleast_1d(scales)
        x = scales[:, None] * omega
        
        H = np.zeros_like(omega)
        H[omega > 0] = 1
        
        with np.errstate(divide='ignore'):
            log_a = np.log(2) + (beta/gamma) * (1+np.log(gamma) - np.log(beta))
            psifs = np.exp(log_a + beta * np.log(np.abs(x)) - np.abs(x)**gamma) * H

        if derivative:
            return 1j * omega * psifs
        
        return psifs
    
    def freq_domain_numba(self, omega, scales, out, *, derivative=False):
        return _morse_freq_domain(omega,
                                  scales,
                                  self.gamma,
                                  self.beta,
                                  out,
                                  derivative=derivative)
    
    def freq_to_scale(self, freqs):
        
        # input should be in normalized radian frequencies
        return self.wp / np.atleast_1d(freqs)
    
    def scale_to_freq(self, scales):
        return self.wp / np.atleast_1d(scales)
    
    def reference_coi(self, *, threshold=1/(np.e**2)):
        base_scale = self.wp / 1
        base_coi = reference_coi(self.freq_domain, base_scale)
        
        return base_scale, base_coi
        
    def coi(self, scales, reference_scale, reference_coi, *, fs=1):
        
        return coi(scales, reference_scale, reference_coi, fs=fs)
    
    @property
    def admissibility_constant(self):
        
        def c_psi(omega):
            
            return self.freq_domain(omega, 1)[0] / omega
        
        return quad(c_psi, 0, np.inf)[0]
    
    @property
    def is_analytic(self):
        return True
        
        
@njit
def _amor_freq_domain(omega, scales, w0, out, *, derivative=False):
    
    H = np.zeros_like(omega)
    H[omega > 0] = 1
    
    for ii in range(scales.shape[0]):
        x = scales[ii] * omega
        out[ii] = 2 * np.exp(-(x - w0)**2/2) * H
        if derivative:
            out[ii] *= 1j * omega
            
    return True
        
        
class AmorWavelet(Wavelet):
    
    def __init__(self, *, w0=7):
    
        self.w0 = w0
        
        
    def freq_domain(self, omega, scales, *, derivative=False):
        w0 = self.w0
        
        scales = np.atleast_1d(scales)
        x = scales[:, None] * omega
        
        H = np.zeros_like(omega)
        H[omega > 0] = 1
        
        psifs = 2 * np.exp(-(x - w0)**2 / 2) * H
        
        if derivative:
            return 1j * omega * psifs
        
        return psifs
    
    def freq_domain_numba(self, omega, scales, out, *, derivative=False):
        return _amor_freq_domain(omega,
                                 scales,
                                 self.w0,
                                 out,
                                 derivative=derivative)
    
    def freq_to_scale(self, freqs):
        
        return self.w0 / freqs
    
    def scale_to_freq(self, scales):
        return self.w0 / np.atleast_1d(scales)
    
    def reference_coi(self, *, threshold=1/(np.e**2)):
        base_scale = self.w0 / 1
        base_coi = reference_coi(self.freq_domain, base_scale)
        
        return base_scale, base_coi
        
    def coi(self, scales, reference_scale, reference_coi, *, fs=1):
        
        return coi(scales, reference_scale, reference_coi, fs=fs)
    
    @property
    def admissibility_constant(self):
        
        def c_psi(omega):
            
            return self.freq_domain(omega, 1)[0] / omega
        
        # sometimes have trouble integrating, so we extend bounds
        # to include negative frequencies. This adds a negligible
        # amount to the final result
        return quad(c_psi, -np.inf, np.inf)[0]
    
    @property
    def is_analytic(self):
        return True
    
@njit
def _bump_freq_domain(omega, scales, mu, sigma, out, *, derivative=False):
    
    for ii in range(scales.shape[0]):
        w = (scales[ii] * omega - mu) / sigma
        tmp = 2 * np.exp( 1 - 1/(1 - w*w) )
        tmp[~(np.abs(w) < 1)] = 0
        tmp[~np.isfinite(tmp)] = 0
        out[ii] = tmp
        if derivative:
            out[ii] *= 1j * omega

    return True
    
    
class BumpWavelet(Wavelet):
    
    def __init__(self, *, mu=5, sigma=0.6):
        
        self.mu = mu
        self.sigma = sigma
        
    def freq_domain(self, omega, scales, *, derivative=False):

        scales = np.atleast_1d(scales)
        
        mu = self.mu
        sigma = self.sigma
        
        w = (scales[:, None] * omega - mu) / sigma
        mask = np.abs(w) < 1
        
        psifs = np.zeros_like(w)
        psifs[mask] = 2 * np.exp( 1 - 1/(1 - w[mask]*w[mask]) )
        
        if derivative:
            return 1j * omega * psifs
        
        return psifs
    
    def freq_domain_numba(self, omega, scales, out, *, derivative=False):
        return _bump_freq_domain(omega, scales, self.mu, self.sigma, out,
                                 derivative=derivative)
    
    def freq_to_scale(self, freqs):
        
        return self.mu / freqs
    
    def scale_to_freq(self, scales):
        return self.mu / np.atleast_1d(scales)
    
    def reference_coi(self, *, threshold=1/(np.e**2)):
        base_scale = self.mu / 1
        base_coi = reference_coi(self.freq_domain, base_scale)
        
        return base_scale, base_coi
        
    def coi(self, scales, reference_scale, reference_coi, *, fs=1):
        
        return coi(scales, reference_scale, reference_coi, fs=fs)
    
    @property
    def admissibility_constant(self):
        
        def c_psi(omega):
            
            return self.freq_domain(omega, 1)[0] / omega
        
        return quad(c_psi, self.mu - self.sigma, self.mu + self.sigma)[0]
    
    @property
    def is_analytic(self):
        return True
```
