## Supplementary material for "GhostiPy: an efficient signal processing and spectral analysis toolbox for large data": Source code: wsst.py

```
import time
import numpy as np
from multiprocessing import cpu_count
from numba import njit, prange #set_num_threads, get_num_threads)
from ghostipy.spectral.cwt import cwt
from ghostipy.spectral.wavelets import (MorseWavelet, AmorWavelet, BumpWavelet)

__all__ = ['wsst']

@njit(parallel=True)
def _assign_to_frequency_bins(W, dW, freq_vec, voices_per_octave, eps, fs, n_threads):
    #n_threads_original = get_num_threads() # only available on 0.49.1? fix
    #set_num_threads(n_threads)
    J, T = W.shape
    sst = np.zeros((freq_vec.shape[0], W.shape[1]), dtype=W.dtype)
    
    #phasetf = (dW / W).imag / (2 * np.pi)
    #freq_inds = np.zeros(W.shape[1], dtype=np.int32)
    phasetf = np.zeros(T)
    k = np.zeros(T, dtype=np.int64)
    
    for j in range(J):
        for m in prange(T):
            if np.abs(W[j, m]) > eps:
                phasetf[m] = (dW[j, m] / W[j, m]).imag / (2 * np.pi) * fs
                if (np.isfinite(phasetf[m])
                and phasetf[m] <= freq_vec[0]
                and phasetf[m] >= freq_vec[-1]):
                    k[m] = np.argmin(np.abs(freq_vec - phasetf[m]))
                    #print(j, m, phasetf, k)
                    sst[k[m], m] += W[j, m] * np.log(2) / voices_per_octave
          
#             freq_ind = J - np.floor(J/(log2f2 - log2f1) * (np.log2(phasetf) - log2f1))
#             if (np.isfinite(freq_ind) == True
#             and freq_ind > -1
#             and freq_ind < J
#             and np.abs(W[j, m]) > eps):
                
#                 sst[int(freq_ind), m] = sst[int(freq_ind), m] + W[j, m] * np.log(2) / voices_per_octave
                
    # only available on 0.49.1? fix
    # set_num_threads(n_threads_original)
    return sst

@njit(cache=True)
def _assign_to_frequency_bins_serial(W, dW, freq_vec, voices_per_octave, eps, fs):
    J, T = W.shape
    sst = np.zeros((freq_vec.shape[0], W.shape[1]), dtype=W.dtype)
    
    #phasetf = (dW / W).imag / (2 * np.pi)
    #freq_inds = np.zeros(W.shape[1], dtype=np.int32)
    
    for j in range(J):
        for m in range(T):
            if np.abs(W[j, m]) > eps:
                phasetf = (dW[j, m] / W[j, m]).imag / (2 * np.pi) * fs
                if (np.isfinite(phasetf)
                    and phasetf <= freq_vec[0]
                    and phasetf >= freq_vec[-1]):

                    k = np.argmin(np.abs(freq_vec - phasetf))
                    #print(j, m, phasetf, k)
                    sst[k, m] += W[j, m] * np.log(2) / voices_per_octave
          
#             freq_ind = J - np.floor(J/(log2f2 - log2f1) * (np.log2(phasetf) - log2f1))
#             if (np.isfinite(freq_ind) == True
#             and freq_ind > -1
#             and freq_ind < J
#             and np.abs(W[j, m]) > eps):
                
#                 sst[int(freq_ind), m] = sst[int(freq_ind), m] + W[j, m] * np.log(2) / voices_per_octave
                
    return sst

def compute_sst_chunk(cwt_computed, dcwt_computed, freqs, voices_per_octave,
                      fs, start, n_time_points, eps, sst_out):

    cwt_chunk = cwt_computed[:, start:start+n_time_points]
    dcwt_chunk = dcwt_computed[:, start:start+n_time_points]
    sst_out[:, start:start + n_time_points] = assign_to_frequency_bins_serial(
        cwt_chunk,
        dcwt_chunk,
        freqs,
        voices_per_octave,
        eps,
        fs)

def wsst(data, *, eps=1e-8, fs=1, timestamps=None, wavelet=MorseWavelet(gamma=3, beta=20),
         freq_limits=None, voices_per_octave=32, n_workers=cpu_count(),
         verbose=False, method='full', remove_mean=False, boundary='mirror',
         coi_threshold=1/(np.e**2), describe_dims=False, cwt_out=None,
         sst_out=None, write_chunk_size='1G'):

    """Computes the wavelet synchrosqueezed transform

    Parameters
    ----------
    data : np.ndarray, with shape (n_timepoints, )
        Data with which to compute the CWT
    eps : float, optional
        Minimum absolute value of the CWT coefficients that are
        considered nonzero.
        Default is 1e-8.
    fs : float
        Sampling rate of the data in Hz.
    timestamps : np.ndarray, with shape (n_timepoints, ) optional
        Timestamps corresponding to the data, in seconds.
        If None, they will be computed automatically based on the
        assumption that all the data are one contiguous block.
    wavelet : ghostipy.wavelet
        Type of wavelet to use for the transform.
        Default is a Morse wavelet with beta=3 and gamma=20.
    freq_limits : list, optional
        List of [lower_bound, upper_bound] for frequencies to use,
        in units of Hz. Note that a reference set of frequencies
        is generated on the shortest segment of data since that
        determines the lowest frequency that can be used. If the
        bounds specified by 'freq_limits' are outside the bounds
        determined by the reference set, 'freq_limits' will be
        adjusted to be within the bounds of the reference set.
    voices_per_octave : int, optional
        Number of wavelet frequencies per octave.
        Default is 32.
    n_workers : integer, optional
        Number of parallel jobs to use.
        Default is the total number of CPUs (which may be virtual).
    verbose : boolean, optional
        Whether to print messages displaying this function's progress.
        Default is False.
    method: {'full', 'ola'}, optional
        Which algorithm to use for computing the CWT. 'ola' will give
        superior performance for long lengths of data.
    derivative: boolean, optional
        Whether to compute the derivative of the CWT.
        Default is False.
    remove_mean: boolean, optional
        Whether to remove the mean of the data before computing the CWT.
        Default is False.
    boundary: {'mirror', 'zeros', 'periodic'}, optional
        To handle boundaries, the data are extended before computing the CWT
        according to the following options:
        'mirror' : Mirror/reflect the data at each end
        'zeros': Add zeros at each end
        'periodic': Treat the data as periodic
        Note that regardless of the boundary method, the CWT should not be
        treated as reliable within the cone of influence.
        Default is 'mirror'.
    coi_threshold : float, optional
        The value C that determines the wavelet's cone of influence. The
        maximum value P of the wavelet's power autocorrelation is taken
        in the time domain. Then the cone of influence is given by the
        region where the power autocorrelation is above C*P. Default value
        for C is e^(-2).
    describe_dims : boolean, optional
        Whether to return the expected shape and dtype of the intermediate
        and final output arrays and return immediately (no SST is computed).
        This option is useful for out-of-core computation. While the expected
        shape should not be changed, the dtype is only suggested, e.g. it is
        acceptable to use a lower precision dtype (i.e. complex64 instead
        of complex128 to save space).
        Default is False.
    cwt_out: array-like, optional
        If specified, the CWT coefficients (used in an intermediate step)
        will be stored here. Useful if there is not enough space in memory
        must instead be saved to an array stored on disk.
    sst_out : array-like, optional
        If specified, the final wavelet SST coefficients will be stored here.
        Useful if the output is too large to fit into memory and must instead
        be saved to an array stored on disk.
    write_chunk_size : string, optional
        Maximum chunk size to use when writing to the final SST output
        array. This option is most useful if the array is stored on disk
        (specifically, too many write calls may result in lower performance).
        Use 'K' for kilobytes, 'M' for megabytes, and 'G' for gigabytes.
        Default is 1 GB.

    Returns
    -------
    If 'describe_dims' is True:
        cwt_shape, cwt_dtype, sst_shape, sst_dtype : tuple
            A tuple consisting of:
            (1) expected shape of the intermediate CWT coefficient array,
            (2) dtype of the intermediate CWT array,
            (3) expected shape of the final output SST coefficient array,
            (4) dtype of the final output SST coefficient array.
    Otherwise:
        coefs : np.ndarray, with shape (n_scales, n_timepoints)
            Calculated continuous wavelet coefficients. Note that the scale
            dimension is ordered by increasing wavelet scale, which corresponds
            to decreasing frequency.
        scales : np.ndarray, with shape (n_scales, )
            Wavelet scales for which CWT was calculated.
        frequencies : np.ndarray, with shape (n_frequencies, )
            If a sampling rate is given, these will be the frequency
            corresponding to each wavelet scale.
        timestamps : np.array, with shape (n_timepoints, )
            If timestamps were not specified, these are the timestamps
            calculated for data samples. Pass through of input 
            timestamps otherwise.
        cois : np.ndarray, with shape (n_cois, )
            Cones of influence for each wavelet scale.
    """

    t0 = time.time()
    
    if describe_dims:
        if verbose:
            print("Calculating output array sizes. Skipping SST transform")
        shape, dtype = cwt(
            data, fs=fs,
            timestamps=timestamps,
            wavelet=wavelet,
            freq_limits=freq_limits, 
            voices_per_octave=voices_per_octave,
            n_workers=n_workers,
            verbose=verbose,
            method=method,
            remove_mean=remove_mean,
            boundary=boundary,
            coi_threshold=coi_threshold,
            describe_dims=describe_dims)

        # same shape and dtype for CWT and SST
        return shape, dtype, shape, dtype

    if verbose:
        print("Computing CWT")
    cwt_computed, scales, freqs, timestamps, cois = cwt(
            data, fs=fs,
            timestamps=timestamps,
            wavelet=wavelet,
            freq_limits=freq_limits, 
            voices_per_octave=voices_per_octave,
            n_workers=n_workers,
            verbose=verbose,
            method=method,
            derivative=False,
            remove_mean=remove_mean,
            boundary=boundary,
            coi_threshold=coi_threshold,
            describe_dims=describe_dims,
            cwt_out=cwt_out)

    if verbose:
        print("Computing the CWT derivative")
    # fill output array with derivative, will be overwritten later with SST data
    dcwt_computed, _, _, _, _ = cwt(
            data, fs=fs,
            timestamps=timestamps,
            wavelet=wavelet,
            freq_limits=freq_limits, 
            voices_per_octave=voices_per_octave,
            n_workers=n_workers,
            verbose=verbose,
            method=method,
            derivative=True,
            remove_mean=remove_mean,
            boundary=boundary,
            coi_threshold=coi_threshold,
            describe_dims=describe_dims,
            cwt_out=sst_out)
    
    # compute sst on chunks
    A, T = dcwt_computed.shape
    n_tot_elements = dcwt_computed.size
    n_bytes_per_element = dcwt_computed.dtype.itemsize

    if write_chunk_size is None:
        n_bytes = n_tot_elements * n_bytes_per_element
    elif write_chunk_size[-1] == 'K':
        n_bytes = int(write_chunk_size[:-1]) * 1000
    elif write_chunk_size[-1] == 'M':
        n_bytes = int(write_chunk_size[:-1]) * 1000**2
    elif write_chunk_size[-1] == 'G':
        n_bytes = int(write_chunk_size[:-1]) * 1000**3
    else:
        raise ValueError("Unsupported value of write_chunk_size")

    if n_bytes > n_tot_elements * n_bytes_per_element:
        n_time_points = T
    else:
        n_time_points = int(np.floor(n_bytes / (A * n_bytes_per_element)))
    starts = np.arange(0, T, n_time_points)

    sst_out = dcwt_computed

    for ii, start in enumerate(starts):
        if verbose:
            print(f"Computing SST chunk {ii} of {len(starts) - 1}")
        
        cwt_chunk = cwt_computed[:, start:start+n_time_points]
        dcwt_chunk = dcwt_computed[:, start:start+n_time_points]
        n_actual_points = cwt_chunk.shape[1]
        sst_out[:, start:start + n_actual_points] = _assign_to_frequency_bins(
            cwt_chunk,
            dcwt_chunk,
            freqs,
            voices_per_octave,
            eps,
            fs,
            n_workers)

    if verbose:
        print(f'WSST total elapsed time: {time.time() - t0} seconds')
        
    return sst_out, scales, freqs, timestamps, cois
```
