## Supplementary material for "GhostiPy: an efficient signal processing and spectral analysis toolbox for large data": Source code: utils.py

```
import numpy as np

__all__ = ['hz_to_normalized_rad',
           'normalized_rad_to_hz']

def hz_to_normalized_rad(freqs, fs):
    return freqs / fs * 2 * np.pi

def normalized_rad_to_hz(rad, fs):
    return rad / np.pi * fs / 2
```
