## Supplementary material for "GhostiPy: an efficient signal processing and spectral analysis toolbox for large data": Source code: README.md

# ghostipy

## Installation
The easiest way to install is ```pip install ghostipy```. You can also install from this repo
using the ```master``` branch

## New users
Example notebooks illustrating common use cases will be uploaded soon. If you have any questions or would like to request a feature, do not hesitate to submit an issue. We are very responsive and would love to provide a solution for your specific needs.

## Branch guide
```master```: The most stable branch and the one end users should prefer by default.

```develop```: Contains features that are still in draft mode and have not been imported into the global ```ghostipy``` namespace. Features are generally expect to work, but not guaranteed.

```unstable```: The main development branch. Not every feature may work; use at your own risk.
