## Supplementary material for "GhostiPy: an efficient signal processing and spectral analysis toolbox for large data": Source code: setup.py

```
from setuptools import setup, find_packages

# long_description = read('README.rst')

def get_version_string():
    version = {}
    with open("ghostipy/version.py") as f:
        exec(f.read(), version)
    return version["__version__"]

setup(
    name='ghostipy',
    version=get_version_string(),
    license='Apache License',
    author='Joshua Chu',
    install_requires=['numpy>=1.18.1',
                      'scipy>=1.4.1',
                      'pyfftw>=0.12.0',
                      'numba>=0.48.0',
                      'dask>=2.15.0',
                      'xarray>=0.15.1',
                      'matplotlib>=3.1.3',
                      'datashader>=0.10.0',
                      'hvplot>=0.5.2'
                     ],
    author_email='',
    description='Signal processing and spectral analysis toolbox',
    packages=find_packages(),
    keywords="neuroscience spectral analysis",
    include_package_data=True,
    platforms='any',
    classifiers=['Programming Language :: Python :: 3']
)
```
