## Supplementary material for "GhostiPy: an efficient signal processing and spectral analysis toolbox for large data": Notebooks for figures: Figure1.ipynb

```
# %% [code]
# You may need to uncomment the following line if running this
# notebook on Colab
# %pip install ghostipy

import ghostipy as gsp
import numpy as np
import matplotlib.pyplot as plt
import h5py
from urllib.request import urlopen

# %% [code]
# Download example data to a local file
s3_url = 'https://ghostipy-example-data.s3-us-west-2.amazonaws.com/example-ripple.h5'
with urlopen(s3_url) as response, open('example-ripple.h5', 'wb') as out_file:
    data = response.read()
    out_file.write(data)

# %% [code]
# Load numpy arrays from example data file
with h5py.File('example-ripple.h5', 'r') as hf:
    ripple_data = hf['ripple_data'][:]
    fs = hf['sampling_rate'][()]

# %% [code]
fig, axes = plt.subplots(2, 2, figsize=(12.5, 8))
fig.text(0.5, 0.03, 'Frequency (Hz)', ha='center', fontsize=22)
fig.text(0.012, 0.5, '(units^2 / Hz)', va='center', fontsize=22, rotation='vertical')

labels = ['a', 'b', 'c', 'd']

for subax_array in axes:
    
    for subax in subax_array:

        for spine in subax.spines:
            subax.spines[spine].set_linewidth(3)
        subax.spines['top'].set_visible(False)
        subax.spines['right'].set_visible(False)

        subax.tick_params(labelsize=20, width=4, length=5)

# show this part
bandwidths = [10, 15, 20, 25]
for ii, bandwidth in enumerate(bandwidths):
    psd, freqs = gsp.mtm_spectrum(ripple_data,
                                  fs=fs,
                                  n_fft_threads=8,
                                  bandwidth=bandwidth)
    row_idx, col_idx = np.unravel_index(ii, (2, 2))
    axes[row_idx, col_idx].plot(freqs, psd, lw=4, color='#064175')

    axes[row_idx, col_idx].set_xlim(50, 500)
    axes[row_idx, col_idx].set_ylim(0, 2000)
    
    axes[row_idx, col_idx].set_title("Bandwidth: {} Hz".format(bandwidths[ii]), fontsize=24)
    axes[row_idx, col_idx].text(-0.25, 1.1, labels[ii], transform=axes[row_idx, col_idx].transAxes, 
                                size=27, weight='bold')
    
plt.subplots_adjust(wspace=0.3, hspace=0.4)

# %% [code]
```
