## Supplementary material for "GhostiPy: an efficient signal processing and spectral analysis toolbox for large data": Notebooks for figures: Figure2.ipynb

```
# %% [code]
# You may need to uncomment the following line if running this
# notebook on Colab
# %pip install ghostipy

import ghostipy as gsp
import numpy as np
import scipy.signal as sig
import matplotlib.pyplot as plt
import h5py
import warnings
from urllib.request import urlopen

# %% [code]
# Download example data to a local file
s3_url = 'https://ghostipy-example-data.s3-us-west-2.amazonaws.com/example-ripple-extended.h5'
with urlopen(s3_url) as response, open('example-ripple-extended.h5', 'wb') as out_file:
    data = response.read()
    out_file.write(data)

# %% [code]
# Load numpy arrays from example data file
with h5py.File('example-ripple-extended.h5', 'r') as hf:
    ripple_data = hf['ripple_data'][:]
    fs = hf['sampling_rate'][()]

# %% [code]
nperseg = 64
noverlap = 16
bw = 25

f_spect, t_spect, psd_spect = sig.spectrogram(
    ripple_data, fs=fs, nperseg=nperseg, noverlap=noverlap)

psd_mtm, f_mtm, t_mtm = gsp.mtm_spectrogram(
    ripple_data, bw, fs=fs, nperseg=nperseg, noverlap=noverlap)

# warnings can be ignored for this particular case
with warnings.catch_warnings():
    warnings.simplefilter("ignore")
    coefs_cwt, _, f_cwt, t_cwt, _ = gsp.cwt(
        ripple_data, fs=fs, freq_limits=[1, 500])

    coefs_wsst, _, f_wsst, t_wsst, _ = gsp.wsst(
        ripple_data, fs=fs, freq_limits=[1, 500],
        voices_per_octave=32)

# will be normalized such that max is 1, so the
# sampling rate factor can be dropped
psd_cwt = coefs_cwt.real**2 + coefs_cwt.imag**2
psd_wsst = coefs_wsst.real**2 + coefs_wsst.imag**2

psd_spect /= np.max(psd_spect)
psd_mtm /= np.max(psd_mtm)
psd_cwt /= np.max(psd_cwt)
psd_wsst /= np.max(psd_wsst)

# %% [code]
fig, axes = plt.subplots(2, 2, figsize=(12.5, 8))

fig.text(0.5, 0.03, 'Frequency (Hz)', ha='center', fontsize=22)
fig.text(0.012, 0.5, 'Normalized PSD', va='center', fontsize=22, rotation='vertical')

labels = ['a', 'b', 'c', 'd']
titles = ['Spectrogram', 'MT Spectrogram', 'CWT Spectrogram', 'SST Spectrogram']
kwargs_dict = {}
kwargs_dict['cmap'] = plt.cm.Spectral_r
kwargs_dict['vmin'] = 0
kwargs_dict['vmax'] = 1
kwargs_dict['linewidth'] = 0
kwargs_dict['rasterized'] = True

for subax_array in axes:
    for subax in subax_array:
        subax.tick_params(labelsize=20, width=3, length=5)
        subax.set_ylim(0, 400)

pc_spect = axes[0, 0].pcolormesh(t_spect, f_spect, psd_spect, **kwargs_dict)
pc_spect.set_edgecolor('face')
cbar_spect = fig.colorbar(pc_spect, ax=axes[0, 0])
cbar_spect.ax.tick_params(labelsize=16)
axes[0, 0].set_title("Scipy Spectrogram", fontsize=24)
axes[0, 0].text(-0.25, 1.1, 'a', transform=axes[0, 0].transAxes, 
                size=27, weight='bold')

pc_mtm = axes[0, 1].pcolormesh(t_mtm, f_mtm, psd_mtm, **kwargs_dict)
pc_mtm.set_edgecolor('face')
cbar_mtm = fig.colorbar(pc_mtm, ax=axes[0, 1])
cbar_mtm.ax.tick_params(labelsize=16)
axes[0, 1].set_title("MT Spectrogram", fontsize=24)
axes[0, 1].text(-0.25, 1.1, 'b', transform=axes[0, 1].transAxes, 
                size=27, weight='bold')

pc_cwt = axes[1, 0].pcolormesh(t_cwt, f_cwt, psd_cwt, **kwargs_dict)
pc_cwt.set_edgecolor('face')
cbar_cwt = fig.colorbar(pc_cwt, ax=axes[1, 0])
cbar_cwt.ax.tick_params(labelsize=16)
axes[1, 0].set_title("CWT Spectrogram", fontsize=24)
axes[1, 0].text(-0.25, 1.1, 'c', transform=axes[1, 0].transAxes, 
                size=27, weight='bold')

pc_wsst = axes[1, 1].pcolormesh(t_wsst, f_wsst, psd_wsst, **kwargs_dict)
pc_wsst.set_edgecolor('face')
cbar_wsst = fig.colorbar(pc_wsst, ax=axes[1, 1])
cbar_wsst.ax.tick_params(labelsize=16)
axes[1, 1].set_title("SST Spectrogram", fontsize=24)
axes[1, 1].text(-0.25, 1.1, 'd', transform=axes[1, 1].transAxes, 
                size=27, weight='bold')

plt.subplots_adjust(wspace=0.3, hspace=0.4)

# %% [code]
```
