## Supplementary material for "GhostiPy: an efficient signal processing and spectral analysis toolbox for large data": Notebooks for figures: Figure3.ipynb

```
# %% [code]
# You may need to uncomment the following line if running this
# notebook on Colab
# %pip install ghostipy

import ghostipy as gsp
import numpy as np
import matplotlib.pyplot as plt
import scipy.signal as sig
import matplotlib.ticker as plticker

# %% [code]
tw = 2
p = 2
fs = 30000

numtaps = gsp.estimate_taps(fs, tw)
band_edges = [4, 6, 10, 12]
desired =    [0, 1,  1,  0]

theta_filter = gsp.firdesign(numtaps, band_edges, desired,
                             fs=fs, p=p)

# %% [code]
numtaps = gsp.estimate_taps(30000, 2)
band_edges = [10, 12, 55, 60, 100, 150, 200, 250, 300, 350]
desired =    [ 1,  0,  0,  1,   1,   0,   0,   1,   1,   0]

arbitrary_filter = gsp.firdesign(numtaps, band_edges, desired, fs=fs, p=p)

# %% [code]
w_theta, h_theta = sig.freqz(theta_filter, worN=65536*16)
w_arbitrary, h_arbitrary = sig.freqz(arbitrary_filter, worN=65536*16)

# %% [code]
import matplotlib.ticker as plticker

w, h = sig.freqz(theta_filter, worN=65536*16)
fig, axes = plt.subplots(1, 2, figsize=(12.5, 4))
ax1, ax2 = axes

for ax in axes:
    for spine in ('left', 'bottom'):
        ax.spines[spine].set_linewidth(3)
    for spine in ('top', 'right'):
        ax.spines[spine].set_visible(False)
    ax.tick_params(labelsize=20, width=3, length=7, axis='both')

ax1.plot(w_theta/np.pi*fs/2, np.abs(h_theta), linewidth=3)
ax1.set_title("Theta band", fontsize=24)
ax1.plot([5, 11], [0.5, 0.5], 'o', color='g', fillstyle='none', markersize=10, markeredgewidth=3)
loc = plticker.MultipleLocator(base=2.0) # this locator puts ticks at regular intervals
ax1.xaxis.set_major_locator(loc)
ax1.tick_params(labelsize=20, width=4, length=7, axis='both')
ax1.text(-0.2, 1.1, 'a', transform=ax1.transAxes, 
        size=27, weight='bold')
ax1.set_xlim(0, 17)

ax2.plot(w_arbitrary/np.pi*fs/2, np.abs(h_arbitrary), linewidth=3)
ax2.set_title("Arbitrary bands", fontsize=24)
ax2.plot([11, 57.5, 125, 225, 325],
         [0.5, 0.5, 0.5, 0.5, 0.5],
         'o', color='g', fillstyle='none',
         markersize=10, markeredgewidth=3)
ax2.tick_params(labelsize=20, width=4, length=7)
ax2.text(-0.2, 1.1, 'b', transform=ax2.transAxes, 
        size=27, weight='bold')
ax2.set_xlim(0, 400)

fig.text(0.5, -0.08, 'Frequency (Hz)', ha='center', fontsize=22)
fig.text(0.03, 0.5, 'Magnitude Response', va='center', fontsize=22, rotation='vertical')

plt.subplots_adjust(wspace=0.3)

# %% [code]
```
