## Supplementary material for "GhostiPy: an efficient signal processing and spectral analysis toolbox for large data": Notebooks for figures: Figure4.ipynb

```
# %% [code]
# You may need to uncomment the following line if running this
# notebook on Colab
# %pip install ghostipy

import numpy as np
import matplotlib.pyplot as plt
import urllib
import ghostipy as gsp
import h5py
from urllib.request import urlopen

# %% [code]
# Download example data to a local file
s3_url = 'https://ghostipy-example-data.s3-us-west-2.amazonaws.com/example_data.h5'
with urlopen(s3_url) as response, open('example_data.h5', 'wb') as out_file:
    data = response.read()
    out_file.write(data)

# %% [code]
# Load numpy arrays from example data file
with h5py.File('example_data.h5', 'r') as hf:
    lfp = hf['lfp'][:]
    lfp_ts = hf['timestamps'][:]
    speed = hf['speed'][:]
    ts_pos = hf['speed_timestamps'][:]

# %% [code]
# Zoom in on a particular window of time
t1 = 274
t2 = 278.5

idx = (lfp_ts >= t1) & (lfp_ts <= t2)

LFP = lfp[idx]
ts = lfp_ts[idx]

idx = (ts_pos >= t1) & (ts_pos <= t2)

spd = speed[idx]
ts_spd = ts_pos[idx]

# %% [code]
fs_lfp = 1500

gamma = 3 
beta = 10 # decrease beta to improve temporal resolution

cwtcoefs, _, freq, cwtts, _ = gsp.cwt(
    LFP, timestamps=ts, freq_limits=[1, 250], fs=fs_lfp,
    wavelet=gsp.MorseWavelet(gamma=gamma, beta=beta))

cwt_power = cwtcoefs.imag**2 + cwtcoefs.real**2

plt.rcParams.update({'axes.spines.right': False, 'axes.spines.top': False, 'pdf.fonttype':42})
fig, axes = plt.subplots(3, 1, figsize=(6, 4), sharex=True, 
                         gridspec_kw={'hspace':0.15, 'height_ratios':[0.1,0.9,0.2]})
ax1, ax2, ax3 = axes

ax1.plot(ts_spd, spd)
ax1.set_ylim(-1, 10)
ax1.set_ylabel("Speed\n(cm/s)", labelpad=10)
ax1.tick_params(axis='x', which='both', bottom=False, top=False)

t_ax, f_ax = np.meshgrid(ts, freq)
ax2.pcolormesh(t_ax, f_ax, cwt_power, shading='gouraud', 
               cmap=plt.cm.viridis, vmin=0, vmax=12500, rasterized=True)
ax2.set_ylabel("Frequency (Hz)")
ax2.set_ylim(1, 250)
ax2.tick_params(axis='x', which='both', bottom=False, top=False)

ax3.plot(ts, LFP, 'grey', linewidth=0.5)

# %% [code]
```
