## Supplementary material for "GhostiPy: an efficient signal processing and spectral analysis toolbox for large data": Notebooks for figures: Figure5.ipynb

```
# %% [code]
# You may need to uncomment the following line if running this
# notebook on Colab
# %pip install ghostipy

import numpy as np
import h5py
import matplotlib.pyplot as plt
from urllib.request import urlopen

# %% [code]
# Download example data to a local file
s3_url = 'https://ghostipy-example-data.s3-us-west-2.amazonaws.com/cwt-comparison.h5'
with urlopen(s3_url) as response, open('cwt-comparison.h5', 'wb') as out_file:
    data = response.read()
    out_file.write(data)

# %% [code]
# Load numpy arrays from example data file
with h5py.File('cwt-comparison.h5', 'r') as hf:
    ks = hf['ks'][:]
    matlab_durations = hf['matlab_durations'][:]
    matlab_max_memory = hf['matlab_max_memory'][:]
    python_durations = hf['python_durations'][:]
    python_max_memory = hf['python_max_memory'][:]

# %% [code]
fig, ax = plt.subplots(1, 2, figsize=(18, 5))

for subax in ax:

    for spine in ('left', 'bottom'):
        subax.spines[spine].set_linewidth(3)
    for spine in ('top', 'right'):
        subax.spines[spine].set_visible(False)

    subax.tick_params(axis='both', labelsize=20, width=3, length=5)

ax[0].plot(ks / 1e6, matlab_durations, color='#e03631', linewidth=3)
ax[0].plot(ks / 1e6, python_durations, color='#1f77b4', linewidth=3)
ax[0].set_xlabel("Data length (Megasamples)", fontsize=20)
ax[0].set_ylabel('Computation time (sec)', fontsize=20)
ax[0].text(-0.22, 1.1, 'a', transform=ax[0].transAxes, 
         size=20, weight='bold')

ax[1].plot(ks / 1e6, matlab_max_memory, color='#e03631', linewidth=3)
ax[1].plot(ks / 1e6, python_max_memory, color='#1f77b4', linewidth=3)
ax[1].set_xlabel("Data length (Megasamples)", fontsize=20)
ax[1].set_ylabel('Peak memory usage (MB)', fontsize=20)
ax[1].text(-0.22, 1.1, 'b', transform=ax[1].transAxes,
         size=20, weight='bold')

leg0 = ax[0].legend(['MATLAB', 'Ghostipy'], prop={'size': 15})
leg1 = ax[1].legend(['MATLAB', 'Ghostipy'], prop={'size': 15})

plt.subplots_adjust(wspace=0.3)

# %% [code]
```
